# Large domain movements through lipid bilayer mediate substrate release and inhibition of glutamate transporters

**DOI:** 10.1101/2020.05.19.103432

**Authors:** Xiaoyu Wang, Olga Boudker

## Abstract

Glutamate transporters are essential players in glutamatergic neurotransmission in the brain, where they maintain extracellular glutamate below cytotoxic levels and allow for rounds of transmission. The structural bases of their function are well established, particularly within a model archaeal homologue, sodium and aspartate symporter Glt_Ph_. However, the mechanism of gating on the cytoplasmic side of the membrane remains ambiguous. We report Cryo-EM structures of Glt_Ph_ reconstituted into nanodiscs, including those structurally constrained in the cytoplasm-facing state and either apo, bound to sodium ions only, substrate, or blockers. The structures show that both substrate translocation and release involve movements of the bulky transport domain through the lipid bilayer. They further reveal a novel mode of inhibitor binding and show how solutes release is coupled to protein conformational changes. Finally, we describe how domain movements are associated with the displacement of bound lipids and significant membrane deformations, highlighting the potential regulatory role of the bilayer.

## MAIN TEXT

Sodium and aspartate symporter Glt_Ph_ is an archaeal homologue of human glutamate transporters, which clear the neurotransmitter glutamate from the synaptic cleft following rounds of neurotransmission (Danbolt, 2001). Glt_Ph_ has served as a model system to uncover the structural and mechanistic features of glutamate transporters (Yernool *et al*., 2004; Boudker *et al*., 2007; Reyes *et al*., 2009; Akyuz *et al*., 2013; Erkens *et al*., 2013; Reyes *et al*., 2013; Verdon *et al*., 2014; Akyuz *et al*., 2015; Hanelt *et al*., 2015; Mcilwain *et al*., 2016; Scopelliti *et al*., 2018). Recently, structural studies on other members of the family, including human variants, have enriched the field and have been mostly consistent with earlier findings on Glt_Ph_ (Canul-Tec *et al*., 2017; Garaeva *et al*., 2018; Yu *et al*., 2019). Collectively, these studies provide what appears to be a nearly complete picture of the structural changes that underlie transport. Briefly, the transporters are homotrimers with each protomer consisting of a centrally located scaffold or trimerization domain and a peripheral transport domain that harbors the L-aspartate (L-asp) and three sodium (Na^+^) ions binding sites. The crucial conformational transition from the outward-facing state (OFS), in which L-asp binding site is near the extracellular solution, into the inward-facing state (IFS), from which the substrate is released into the cytoplasm, involves a rigid-body “elevator-like” movement of the transport domain by ca 15 Å across the lipid membrane (Reyes *et al*., 2009; Akyuz *et al*., 2013; Erkens *et al*., 2013; Ruan *et al*., 2017). The structures of the apo transporters in the OFS and IFS showed similar positions of the transport domains that have undergone local structural rearrangements associated with the release of the bound L-asp and Na^+^ ions (Jensen *et al*., 2013; Verdon *et al*., 2014).

The OFS and IFS conformations show a remarkable internal symmetry (Yernool *et al*., 2004; Crisman *et al*., 2009; Reyes *et al*., 2009). In particular, the transport domains feature two pseudo-symmetric helical hairpins (HP) 1 and 2. HP1 lines the interface between the transport and scaffold domains in the OFS, reaching from the cytoplasmic side of the transporter. HP2 lies on the surface of a large extracellular bowl formed by the transporter and occludes L-asp and three Na^+^-binding sites (NA1, 2, and 3). The two hairpins meet near the middle of the lipid bilayer, and their non-helical tips provide essential coordinating moieties for the bound L-asp. As the transport domain translocates into the IFS, HP2 replaces HP1 on the domains interface, while HP1 now lines an intracellular vestibule leading to the substrate-binding site (**Figure 1 Supplementary Figure 1**). Structural and biophysical studies have established that HP2 serves as the extracellular gate of the transporter (Boudker *et al*., 2007; Focke *et al*., 2011; Verdon *et al*., 2014; Riederer e Valiyaveetil, 2019). HP2 closes when the transporter is bound to Na^+^ ions and L-asp and when it is empty (Yernool *et al*., 2004; Jensen *et al*., 2013; Verdon *et al*., 2014). In contrast, it assumes open conformations when the transporter is bound only to Na^+^ ions or Na^+^ ions and competitive blockers DL-*threo-β*-benzyloxyaspartate (TBOA) or (2S,3S)-3-[3-[4- (trifluoromethyl)benzoylamino]benzyloxy]aspartate (TFB-TBOA) (Boudker *et al*., 2007; Verdon *et al*., 2014; Canul-Tec *et al*., 2017).

The gating process in the IFS is less well understood. Based on symmetry considerations, it was first proposed that HP1 might serve as the intracellular gate (Yernool *et al*., 2004) or that the very tip of HP2 might open to release the substrate and ions (Dechancie *et al*., 2011). A large opening of HP2 seemed unlikely because of the steric constraints on the domain interface. However, later structures of a gain-of-function mutant of Glt_Ph_ and human homologous neutral amino acid transporter ASCT2 showed that the transport domain in the IFS could swing away from the scaffold, opening a crevice between the domains (Akyuz *et al*., 2013; Garaeva *et al*., 2018). In this so-called “unlocked” conformation, there was sufficient space for HP2 to open. More recent studies of ASCT2 and of an archaeal GltTk further showed that HP2 could open, suggesting that it serves as a gate in both the OFS and IFS (Garaeva *et al*., 2019; Arkhipova *et al*., 2020). Here, we report a series of Cryo-EM structures of Glt_Ph_ reconstituted into nanodiscs in the IFS and OFS. We show that the transport domain explores a large range of motions in the IFS to which the bilayer adapts through significant bending. These motions are coupled to local changes in HP2 to mediate variable substrate-binding sites exposures to the solvent and accommodate ligands of diverse sizes. They also affect the area of the hydrophobic interface between the transport and scaffold domains. When the transporter is bound to non-transportable blockers or Na^+^ ions only, the area is significantly larger than when the transporter is apo or fully loaded with the substrate and ions. The more extensive interface may contribute to the inability of the transport domains to return to the OFS, providing a mechanism of inhibition and coupled transport.

## RESULTS

### Large range of motions of the transport domain in the IFS

In the outward-facing Glt_Ph_ and EAAT1 in complex with blockers TBOA and TFB-TBOA or Na^+^ ions only, HP2 opens to various degrees, enabling access to the substrate-binding site (Boudker *et al*., 2007; Verdon *et al*., 2014; Canul-Tec *et al*., 2017). To picture gating in the IFS, we imaged the Glt_Ph_ reconstituted into MSP1E3 nanodiscs in the presence of various ligands by single-particle Cryo-EM. We used a variant of Glt_Ph_, conformationally constrained in the IFS by cross-linking of cysteine residues placed into the transport and scaffold domains, Glt_Ph_-K55C/A364C (Glt_Ph_^IFS^) (Reyes *et al*., 2009). Earlier crystal structures of Glt_Ph_^IFS^ pictured the position of the transport domain that was very similar to those visualized in unconstrained inward-facing Glt_Ph_ mutants (Verdon e Boudker, 2012; Akyuz *et al*., 2015).

We determined the structures of Glt_Ph_^IFS^ free of ligands (Glt_Ph_ ^IFS^-Apo-open) or in complex with Na^+^ ions (Glt_Ph_^IFS^-Na) and bound to L-asp (Glt_Ph_^IFS^-Asp), TBOA (Glt_Ph_^IFS^-TBOA), TFB-TBOA (Glt_Ph_^IFS^-TFB-TBOA), and the wide-type outward-facing Glt_Ph_ in complex with TBOA (Glt_Ph_^OFS^-TBOA) to 3.52, 3.66, 3.05, 3.66, 3.71, and 3.66 Å resolution, respectively (**Methods, Figure 1 Supplementary Figures 2–4, and Table 1**). The Glt_Ph_^IFS^-Asp structure was nearly identical to the earlier crystal structure (Reyes *et al*., 2009). The transport domain was well packed against the scaffold primarily through interactions of HP2 and the extracellular part of TM8 (TM8a) with the scaffold TMs 2, 4, and 5. The central axis of the roughly cylindrical transport domain formed a ~35 ° angle with the membrane normal (**Figure 1a, b**). HP2 was closed over the substrate-binding site, and packing between the transport and scaffold domain left no space for the hairpin to open. A similar inter-domain orientation and packing were also observed in a crystal structure of the occluded apo Glt_Ph_^IFS^ (Glt_Ph_^IFS^-Apo-closed, PDB code 4P19, **Figure.1a**). In the new structures of Glt_Ph_^IFS^-Na, Glt_Ph_^IFS^-Apo-open, Glt_Ph_^IFS^-TBOA, and Glt_Ph_^IFS^-TFB-TBOA, approximately the same regions of HP2 and TM8a remained juxtaposed against the scaffold. However, the bulk of the transport domain swung out away from HP2 and the scaffold to different extents (**Figure 1a**) with the largest angle between the transport domain and the membrane normal of ~47° in Glt_Ph_^IFS^-TFB-TBOA (**Figure 1b**). Together, the crystal and Cryo-EM structures define mechanisms of gating in Glt_Ph_ on the extracellular and cytoplasmic sides of the membrane (**Figure 1c, Movie 1**). In the OFS, the bulk of the transport domain remains mostly static relative to the scaffold, and the labile HP2 serves as the extracellular gate. In the IFS, HP2 can maintain interactions with the scaffold, while the bulk of the transport domain swings away to allow access to the binding site. Notably, in a crystal structure of a gain-of-function L-asp-bound mutant Glt_Ph_^IFS^-R276S/M395R, the transport domain is positioned at ~45 ° angle, similar to the Glt_Ph_ ^IFS^-TFB-TBOA (Akyuz *et al*., 2015). However, HP2 remains closed over the binding site and a large lipid-filled gap forms between the transport and scaffold domains. It is currently unclear whether the transport domain first swings away from the scaffold providing space for the consequent HP2 opening or whether HP2 remains in place while the bulk of the domain swings out in a “wag-the-dog” manner.

**Table 1.**
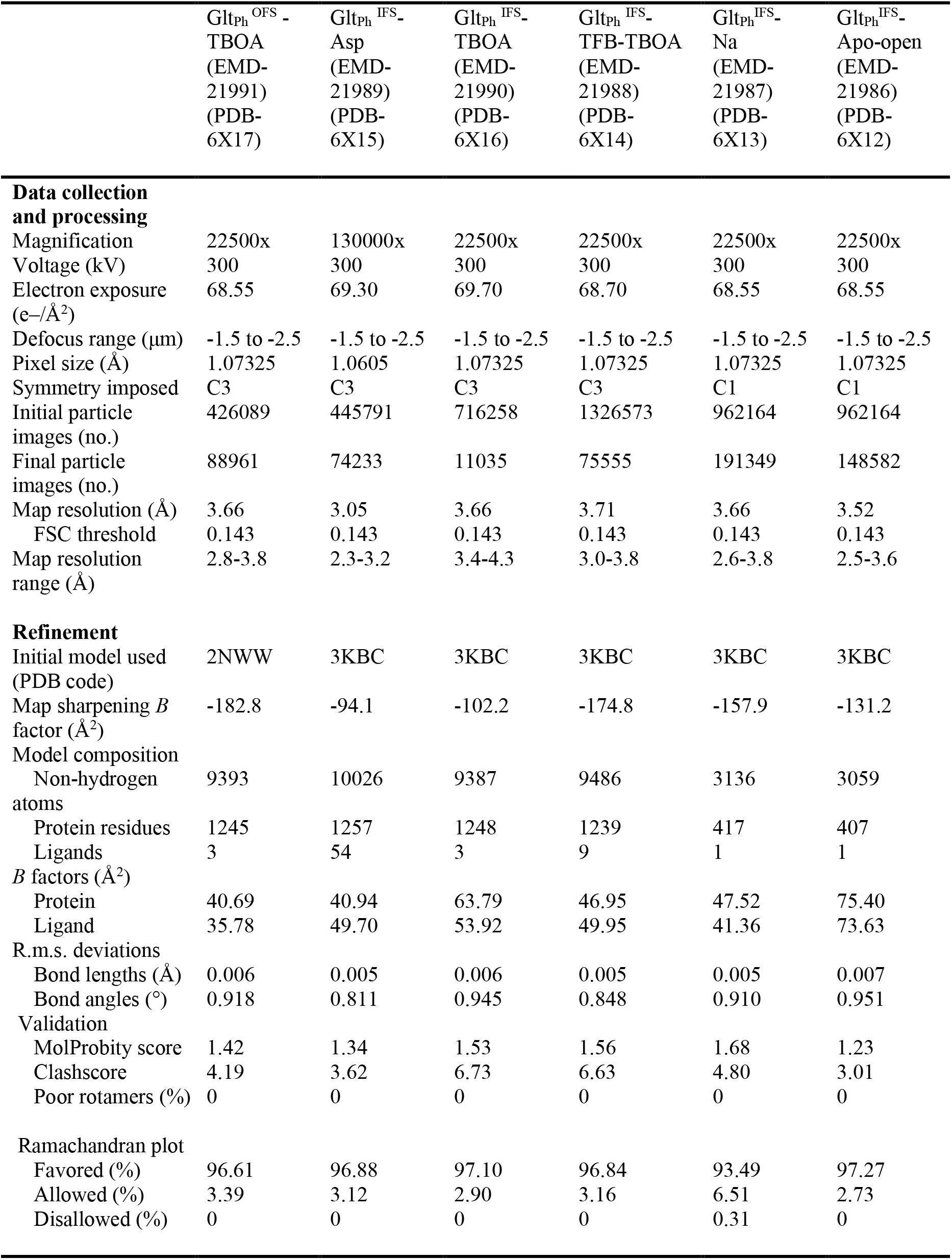
Cryo-EM data collection, refinement and validation statistics

|  | Glt<sub>ph</sub><sup>OFS</sup> -<br>TBOA<br>(EMD-21991)<br>(PDB-6X17) | Glt<sub>ph</sub><sup>IFS</sup> -<br>Asp<br>(EMD-21989)<br>(PDB-6X15) | Glt<sub>ph</sub><sup>IFS</sup> -<br>TBOA<br>(EMD-21990)<br>(PDB-6X16) | Glt<sub>ph</sub><sup>IFS</sup> -<br>TFB-TBOA<br>(EMD-21988)<br>(PDB-6X14) | Glt<sub>ph</sub><sup>IFS</sup> -<br>Na<br>(EMD-21987)<br>(PDB-6X13) | Glt<sub>ph</sub><sup>IFS</sup> -<br>Apo-open<br>(EMD-21986)<br>(PDB-6X12) |
| --- | --- | --- | --- | --- | --- | --- |
| **Data collection and processing** |  |  |  |  |  |  |
| Magnification | 22500x | 130000x | 22500x | 22500x | 22500x | 22500x |
| Voltage (kV) | 300 | 300 | 300 | 300 | 300 | 300 |
| Electron exposure (e-/Å<sup>2</sup>) | 68.55 | 69.30 | 69.70 | 68.70 | 68.55 | 68.55 |
| Defocus range (μm) | -1.5 to -2.5 | -1.5 to -2.5 | -1.5 to -2.5 | -1.5 to -2.5 | -1.5 to -2.5 | -1.5 to -2.5 |
| Pixel size (Å) | 1.07325 | 1.0605 | 1.07325 | 1.07325 | 1.07325 | 1.07325 |
| Symmetry imposed | C3 | C3 | C3 | C3 | C1 | C1 |
| Initial particle images (no.) | 426089 | 445791 | 716258 | 1326573 | 962164 | 962164 |
| Final particle images (no.) | 88961 | 74233 | 11035 | 75555 | 191349 | 148582 |
| Map resolution (Å) | 3.66 | 3.05 | 3.66 | 3.71 | 3.66 | 3.52 |
| FSC threshold | 0.143 | 0.143 | 0.143 | 0.143 | 0.143 | 0.143 |
| Map resolution range (Å) | 2.8-3.8 | 2.3-3.2 | 3.4-4.3 | 3.0-3.8 | 2.6-3.8 | 2.5-3.6 |
| **Refinement** |  |  |  |  |  |  |
| Initial model used (PDB code) | 2NWW | 3KBC | 3KBC | 3KBC | 3KBC | 3KBC |
| Map sharpening *B* factor (Å<sup>2</sup>) | -182.8 | -94.1 | -102.2 | -174.8 | -157.9 | -131.2 |
| Model composition |  |  |  |  |  |  |
| Non-hydrogen atoms | 9393 | 10026 | 9387 | 9486 | 3136 | 3059 |
| Protein residues | 1245 | 1257 | 1248 | 1239 | 417 | 407 |
| Ligands | 3 | 54 | 3 | 9 | 1 | 1 |
| *B* factors (Å<sup>2</sup>) |  |  |  |  |  |  |
| Protein | 40.69 | 40.94 | 63.79 | 46.95 | 47.52 | 75.40 |
| Ligand | 35.78 | 49.70 | 53.92 | 49.95 | 41.36 | 73.63 |
| R.m.s. deviations |  |  |  |  |  |  |
| Bond lengths (Å) | 0.006 | 0.005 | 0.006 | 0.005 | 0.005 | 0.007 |
| Bond angles (°) | 0.918 | 0.811 | 0.945 | 0.848 | 0.910 | 0.951 |
| Validation |  |  |  |  |  |  |
| MolProbity score | 1.42 | 1.34 | 1.53 | 1.56 | 1.68 | 1.23 |
| Clashscore | 4.19 | 3.62 | 6.73 | 6.63 | 4.80 | 3.01 |
| Poor rotamers (%) | 0 | 0 | 0 | 0 | 0 | 0 |
| Ramachandran plot |  |  |  |  |  |  |
| Favored (%) | 96.61 | 96.88 | 97.10 | 96.84 | 93.49 | 97.27 |
| Allowed (%) | 3.39 | 3.12 | 2.90 | 3.16 | 6.51 | 2.73 |
| Disallowed (%) | 0 | 0 | 0 | 0 | 0.31 | 0 |

**Figure 1.**
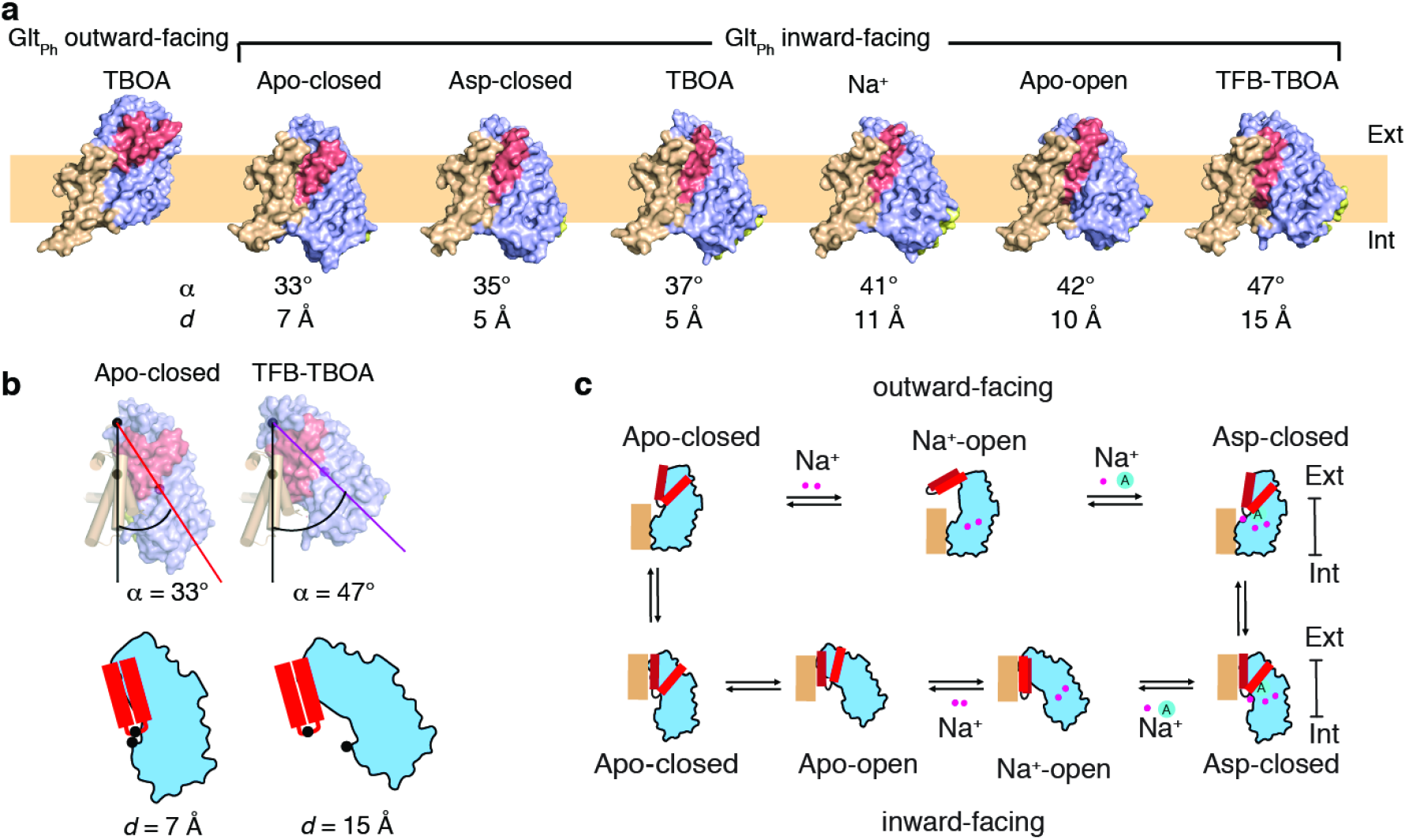
Gating mechanism in the inward-facing state. **a**, Structures of Glt_Ph_ protomers are shown in surface representation viewed in the membrane plain. The scaffold domain is colored wheat, the transport domain blue, HP1 yellow, and HP2 red. The PDB accession code for Glt_Ph_^IFS^-Apo-closed is 4P19. An approximate position of the bilayer is shown as a pale orange rectangle. **b,** Angles between the membrane normal drawn through the center of the scaffold domain and the central axis of the transport domains are shown for Glt_Ph_ ^IFS^-Apo-closed and Glt_Ph_^IFS^-TFB-TBOA (top). Distances between the cα atoms (black circles) of residues R276 and P356 are shown for the same structures under the schematic depiction of the transport domains. Corresponding angles and distances are listed under all structures in panel a. **c,** A schematic representation of the gating mechanism on the extracellular (top) and intracellular (bottom) sides of the membrane.

### Two transporter blockers bind differently to Glt_Ph_^IFS^

TBOA and TFB-TBOA blockers share the amino acid backbone with L-asp but are decorated on β-carbon with one and two bulky benzyl rings, respectively, that cannot fit within the confines of the substrate-binding site. They block transport by binding to the outward-facing Glt_Ph_ or EAATs and arresting HP2 in an open conformation (Boudker *et al*., 2007; Canul-Tec *et al*., 2017). Our Cryo-EM structure of the outward-facing Glt_Ph_^OFS^-TBOA in nanodisc confirmed that the transporter took the same conformation in the absence of crystal contacts in a lipid bilayer (**Figure 2 Supplementary Figure 1a)**. Because TBOA also binds to the IFS of Glt_Ph_ (Reyes *et al*., 2013; Oh e Boudker, 2018), we determined the structures of the Glt_Ph_^IFS^ complexes with the blockers. In the Glt_Ph_^IFS^-TBOA structure, the ligand density was resolved for the aspartate moiety. However, the density for the benzyl group was lacking, perhaps because the blocker is a mixture of L and D isomers leading to heterogeneity (**Figure 2 Supplementary Figure 1b**).

In contrast, in the Glt_Ph_^IFS^-TFB-TBOA structure, TFB-TBOA density was clear, and we modeled the inhibitor in its binding site (**Figure 2a**). We also modeled L-asp into the excess density in the binding site of Glt_Ph_^IFS^-Asp (**Figure 2 Supplementary Figure 1c)**. The bound L-asp and TFB-TBOA share some critical interactions (**Figure 2b)**. Thus, R397 coordinates the side chain carboxylates of aspartate moieties, and D394 coordinates the amino groups. However, TFB-TBOA assumes a different rotomer, leading to the displacement of the backbone carboxylate and the loss of coordination by the highly conserved N401. The aromatic rings of TFB-TBOA protrude from the ligand-binding site and lodge in between the transport and scaffold domains (**Figure 2a, Figure 2 Supplementary Figure 1e**). Most strikingly, HP2 takes a wide-open conformation that is essentially the same as in the outward-facing Glt_Ph_-TBOA complex (**Figure 2 Supplementary Figure 1d**). Interestingly, HP2 was in the same conformation also in an R397C Glt_Ph_ mutant bound to glutamine or benzyl-cysteine. In these structures, the ligands made virtually no interactions with the hairpin but introduced steric clashes disallowing the loop closure (Scopelliti *et al*., 2018). Therefore, it appears that the hairpin intrinsically favors this open conformation.

**Figure 2.**
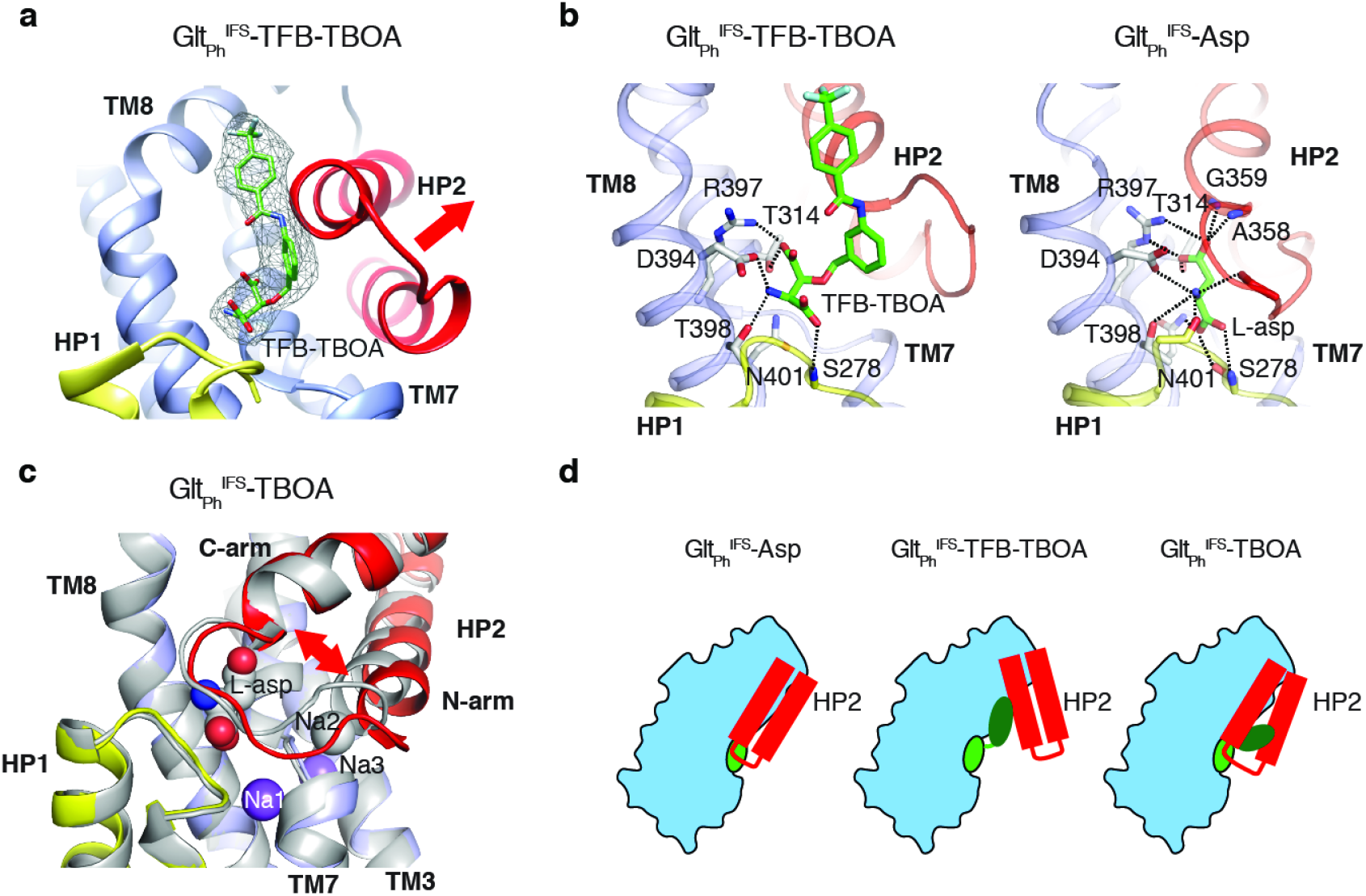
Two mechanisms of blocker binding. **a**, Close-up view of the substrate-binding pocket of Glt_Ph_^IFS^ with bound TFB-TBOA shown in stick representation and colored by atom type. The corresponding density is shown as a black mesh object. Red arrow emphasizes HP2 opining. **b,** TFB-TBOA assumes a distinct rotomer and is coordinated differently from L-asp. **c,** Superimposed Glt_Ph_^IFS^ transport domains in complex with L-asp (grey) and TBOA (colored). Red arrow emphasized parting of the N- and C-terminal arms of HP2. L-asp and Na^+^ ions are shown as spheres. **d**, Two mechanisms of blockers binding to Glt_Ph_^IFS^ showing either opening of HP2 or parting of the two arms to accommodate the bulky moieties of the blockers.

Surprisingly, HP2 does not open in the same way in Glt_Ph_^IFS^-TBOA. Instead, the hairpin remains mostly closed and its N- and C-terminal arms part, perhaps to provide space for the benzyl group (**Figure 2c**). This movement disrupts the Na2 binding site, consistent with previous observations that binding of TBOA and related L-β-threo-benzyl-aspartate to the IFS of the transporter required only two Na^+^ ions (Reyes *et al*., 2013). The movement creates a small opening into the cytoplasmic milieu between the tips of HP1 and HP2. It is not clear whether this conformation reflects a functional state. Perhaps, it recapitulates a transient transporter state, in which a Na^+^ ion has already left the Na2 site while the substrate and two remaining Na^+^ ions are still bound. Water might use the cytoplasmic opening to reach and eventually displace the remaining solutes.

Collectively, the structures show that in Glt_Ph_^IFS^, bulky competitive inhibitors can be accommodated either by opening HP2 or by parting its N- and C-terminal arms (**Figure 2d**). Since the OFS and IFS share the same binding pocket for the substrate and competitive inhibitors, it is likely that the new mode of inhibitor binding, which involves parting of the HP2 arms, can be sampled in the OFS as well. The novel mode of blocker binding might provide new pharmacological avenues for the inhibition of human glutamate transporters.

### M311 and R397 couple HP2 gating to ion and substrate binding

To further explore the gating mechanism, we aimed to resolve a structure of Na^+^ only-bound Glt_Ph_^IFS^ and imaged nanodisc-reconstituted Glt_Ph_^IFS^ frozen in the presence of 200 mM NaCl (**Figure 1 Supplementary Figure 3b and 4, and Table 1**). We isolated two distinct structural classes of Glt_Ph_^IFS^ protomers after symmetry expansion and classification without alignment. The structural heterogeneity was not surprising in retrospect because Na^+^ concentration in the sample was close to the dissociation constant measured for Glt_Ph_^IFS^ (Reyes *et al*., 2013). Thus, we observed both Na^+^-bound and apo states of the transporter, Glt_Ph_^IFS^-Na, and Glt_Ph_^IFS^-Apo-open, respectively. The states were assigned based on the conformations of the conserved non-helical NMD motif (residues 310-312) in TM7, which coordinates of Na^+^ ions in the Na1 and Na3 sites, and TM3, part of the Na3 site (**Figure 3 Supplementary Figure 1a**). In particular, the side chain of M311 protrudes towards the L-asp and Na2 sites in Glt_Ph_^IFS^-Na and Glt_Ph_^IFS^-Asp structures. In contrast, it flips out toward TM3 in our Glt_Ph_^IFS^-Apo-open structure and a published Glt_Ph_^IFS^-Apo-closed crystal structure (Verdon *et al*., 2014). We did not observe density for Na^+^ ions in the Na1 and Na3 sites of Glt_Ph_^IFS^-Na. However, all ion-coordinating residues are positioned similarly to Glt_Ph_^IFS^-Asp (**Figure 3 Supplementary Figure 1b**). Notably, Na1 is coordinated in Glt_Ph_^IFS^-Asp, in part, by an occluded water molecule. In Glt_Ph_^IFS^-Na, the water is no longer occluded and is part of an aqueous cavity (**Figure 3a**). We conclude that ions likely occupy Na1 and Na3 sites, but the Na1 site might be in rapid equilibrium with the solution.

**Figure 3.**
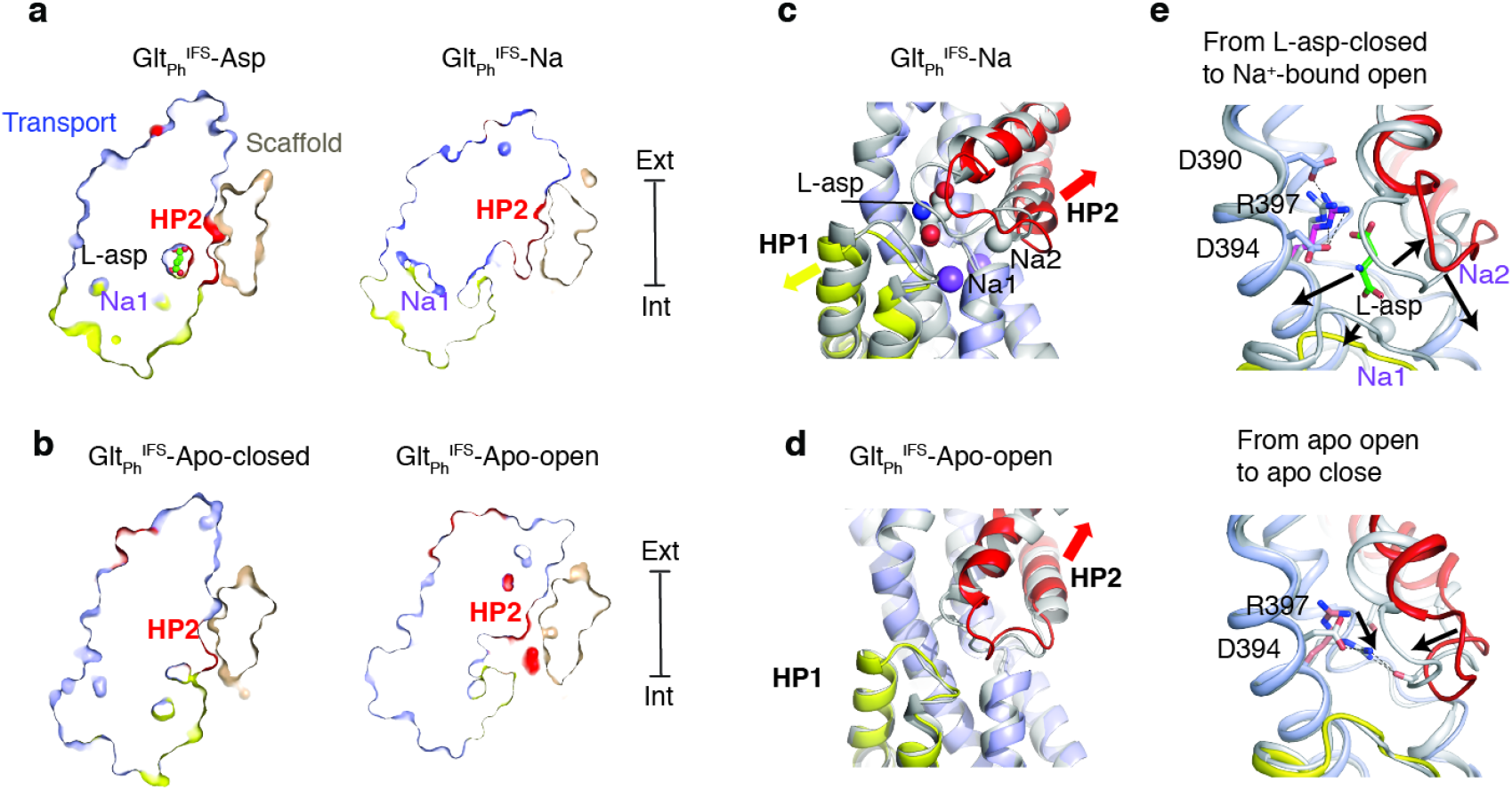
Solute-coupled gating. **a, b,** Thin cross-sections of the protomers taken approximately through L-asp binding sites normal to the membrane plain. The binding site is occluded in Na^+^/L-asp-bound, and closed apo (PDB 4P19) states and is exposed to the solvent in Na^+^-only, and apo open states. **c,** Superimposed transport domains of Glt_Ph_^IFS^-Na (colored) and Glt_Ph_^IFS^-Asp (grey). L-asp and Na^+^ ions are shown as spheres. Yellow and red arrows indicate movements of HP1 and HP2, respectively. **d,** Superimposed transport domains of Glt_Ph_^IFS^-Apo-open (colored) and Glt_Ph_^IFS^-Apo-closed (grey). **e,** Gating steps in the inward-facing state. Top: Local structural changes upon L-asp and Na2 release from Glt_Ph_^IFS^-Asp (grey), leading to an open state of Glt_Ph_^IFS^-Na (colored). Black arrows indicate the release of L-asp and Na2 and associated opening of HP1 and HP2. Bottom: Binding site occlusion transition from Glt_Ph_^IFS^-Apo-open (colored) to Glt_Ph_^IFS^-Apo-closed (grey). Black arrows mark movements of R397 into the binding site and the concomitant closure of HP2.

Surprisingly, Glt_Ph_^IFS^-Apo-open differs significantly from the earlier occluded crystal structure Glt_Ph_ ^IFS^-Apo-closed in that the substrate-binding site is open and hydrated. The opening resembles that in Glt_Ph_^IFS^-Na compared to the occluded Glt_Ph_^IFS^-Asp (**Figure 3a, b**) and shares the overall mechanism: HP2 remains in contact with the scaffold while the rest of the transport domain swings out (**Figure 1c**). From the viewpoint of the transport domain, the conformational changes lead to a similar HP2 opening (**Figure 3c, d, Figure 3 Supplementary Figure 2a**). Interestingly, in Glt_Ph_^IFS^-Na, there is also a small shift of HP1 away from the substrate-binding site, possibly increasing water access to the Na1 site. A similar small movement of the otherwise rigid HP1 was observed in the crystals of apo Glt_Ph_^IFS^ grown in an alkali-free buffer (Verdon *et al*., 2014).

Two residues in the transport domain - M311, and R397 - move significantly during gating and might couple solute binding and release to the large-scale conformational changes. Here we consider a sequence of structural events, which might underlie ion and substrate release in the IFS (**Figure 1c**), starting with Glt_Ph_^IFS^-Asp and going to Glt_Ph_^IFS^-Na, Glt_Ph_^IFS^-Apo-open, and Glt_Ph_^IFS^-Apo-closed (**Movie 2**). In Glt_Ph_^IFS^-Asp, the R397 side chain extends upward, toward the extracellular side of the membrane, where D390 coordinates its guanidinium group. Thus positioned, R397 makes space for L-asp and coordinates its sidechain carboxylate, while D394 coordinates its amino group (**Figure 3e**). M311 protrudes into the binding site and coordinates Na2 (**Figure 3 Supplementary Figure 2**). Extensive interaction of HP2 with the bound L-asp and Na2 likely favor the closed conformation. Upon the release of L-asp and Na2 (Glt_Ph_^IFS^-Na), HP2 opens. R397 is now clamped between D390 and D394, while M311 remains in place (**Figure 3e**, **Figure 3 Supplementary Figure 2b**). The consequent release of Na1 and Na3 leads to a restructuring of the NMD motif and a large outward rotation of M311, which now packs against the open HP2 of Glt_Ph_^IFS^-Apo-open (**Figure 3 Supplementary Figure 2b)**. The guanidinium group of R397 remains between D390 and D394. To achieve the closed apo state, M311 swigs further out into the lipid bilayer, allowing HP2 to close. R397 descends deep into the binding pocket, coordinated now only by D394, and is poised to make direct or through-water interactions with carbonyl oxygens of the closed tip of HP2. Steric hindrance of M311 and more positive local electrostatics may prevent R397 from entering the L-asp binding site, and closing HP2 in Na^+^-only bound Glt_Ph_^IFS^. Physiologically, such Na^+^-bound occluded states should be avoided to prevent Na^+^ leaks.

Interestingly, in our Cryo-EM analysis, we did not find any Glt_Ph_^IFS^-Apo-closed structures previously visualized by crystallography. It might be that the open conformation of the apo Glt_Ph_^IFS^ is the preferred state of the transporter and that the Glt_Ph_^IFS^-Apo-closed state is assumed only transiently, before the outward transition of the transport domain. Packing crystal contacts might have stabilized the closed conformation.

### Ligand-dependent domain interface

HP2 and TM8a comprise most of the transport domain surface interacting with the scaffold. Strikingly, in each of our IFS structures, HP2 takes a different conformation (**Figure 4 Supplementary Figure 1a**). These are similar in structures with Na^+^ ions bound to Na1 and Na3, i.e., in complexes with Na^+^ ions only and with L-asp, TBOA, or TFB-TBOA. The differences are mostly around the tips of HP2 near the L-asp and Na2 sites (**Figure 4 Supplementary Figure 1a and b**). In contrast, the helices restructure significantly in the apo conformations, particularly in Glt_Ph_^IFS^-Apo-open (**Figure 4 Supplementary Figure 1a and c**). Strikingly, when we superimposed all IFS structures, aligning them on the scaffold domain, we observed that the HP2/TM8a motifs present the same bulky hydrophobic residues flanking the flexible tips for interactions with the scaffold: L347, I361, and L378 form virtually the same spatial arrangement. Only in Glt_Ph_^IFS^-Apo-open, where the HP2/TM8a motif, and particularly the N-terminal arm of HP2, is pulled up, I350 replaces L347 (**Figure 4 Supplementary Figure 1d and e**).

Thus, the positions of HP2 tips on the domain interface are mostly conserved. The structural differences in the hairpins then lead to their different orientations relative to the scaffold and different positions of the transport domains, which lean away from the scaffold and rotate around a membrane normal. The rotation is small for Glt_Ph_^IFS^-Apo-closed, relative to Glt_Ph_^IFS^-Asp (7°), but is significant for Glt_Ph_^IFS^-Na (23°), and Glt_Ph_^IFS^-TFB-TBOA (29°). A consequence of these differences is that the bulky residues in the HP2 N-terminal arm, L339, L343, L347, and I350 make more extensive interactions with the scaffold TMs 4a and 4c in Glt_Ph_^IFS^-Na and Glt_Ph_^IFS^-TFB-TBOA compared to other structures. Furthermore, interaction areas between HP2/TM8 and the scaffold domain differ, with Glt_Ph_^IFS^-Apo-closed and Glt_Ph_^IFS^-Asp structures having the smallest area of 1086 and 1076 Å^2^, respectively, and Glt_Ph_^IFS^-Na showing the largest increase of ~400 Å^2^ (**Figure 4**).

**Figure 4.**
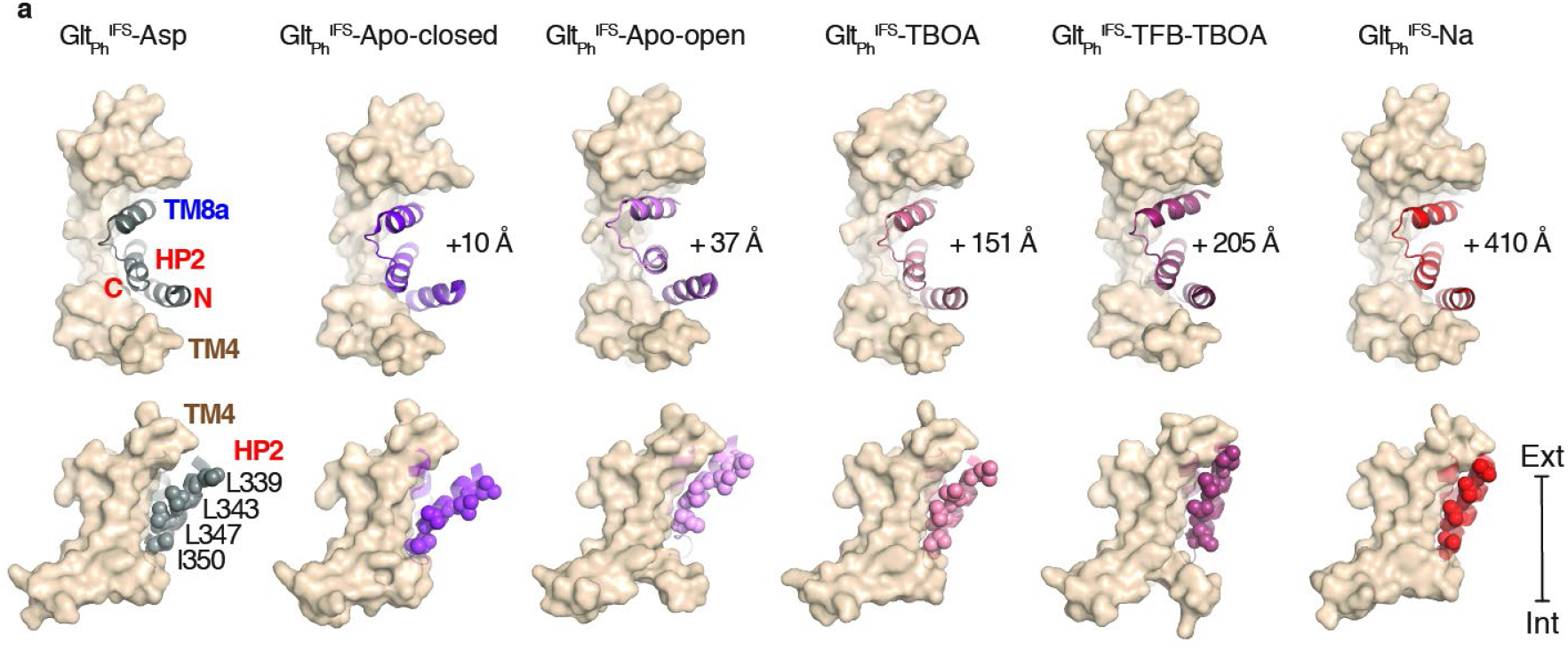
Transport-deficient states show more extensive inter-domain interfaces. **a,** Surface representations of the scaffold domain in light brown, and cartoon representation of HP2/TM8a motif with Glt_Ph_^IFS^-Asp colored grey, Glt_Ph_^IFS^-Apo-closed pink, Glt_Ph_^IFS^-Apo-open purple, Glt_Ph_^IFS^-TBOA salmon, Glt_Ph_^IFS^-TFB-TBOA berry, Glt_Ph_^IFS^-Na red. Side-chains of L339, L343, L347, and L350 are shown as spheres. Upper row: viewed from the extracellular space. The increases in the interdomain interaction surface area relative to Glt_Ph_^IFS^-Asp are shown next to the structures. Lower row: viewed in the membrane plain.

The disruption of the interdomain interface is a prerequisite for the transport domain translocation from the inward-to the outward-facing position. Therefore, altered geometry of the interface and larger interaction area may explain why translocation is inhibited by blockers TBOA, and TFB-TBOA, or especially in the transport domain bound to Na^+^ ions only. While it is not possible to translate interaction areas into energies, it is notable that translocation-competent closed apo and L-asp-bound states show the smallest areas. Consistently, the gain of function mutant R276S/M395R shows a reduced interaction area of only 543 Å^2^ and a translocation rate several-fold faster than the wild type transporter (Akyuz *et al*., 2015).

### Transport domain movements coupled to lipid bilayer

The Cryo-EM structures of the outward- and inward-facing states of Glt_Ph_ are overall similar to the crystal structures. However, they differ in the N-terminus, which is unstructured in crystals but forms a short amphipathic helix positioned on the surface of the nanodiscs in Cryo-EM OFS and IFS structures (**Figure 2 Supplementary Figure 1**). A similar helix was also observed in crystallized EAAT1(Canul-Tec *et al*., 2017). We find highly ordered lipid molecules between the N-terminal helix and the rest of the scaffold at positions conserved in all structures (LipidIn, **Figure 5a and Figure 5 Supplementary Figure 1**). It seems likely that the helix anchors the scaffold domain in the lipid membrane and forms lipid-mediated interactions with the neighboring subunit.

**Figure 5.**
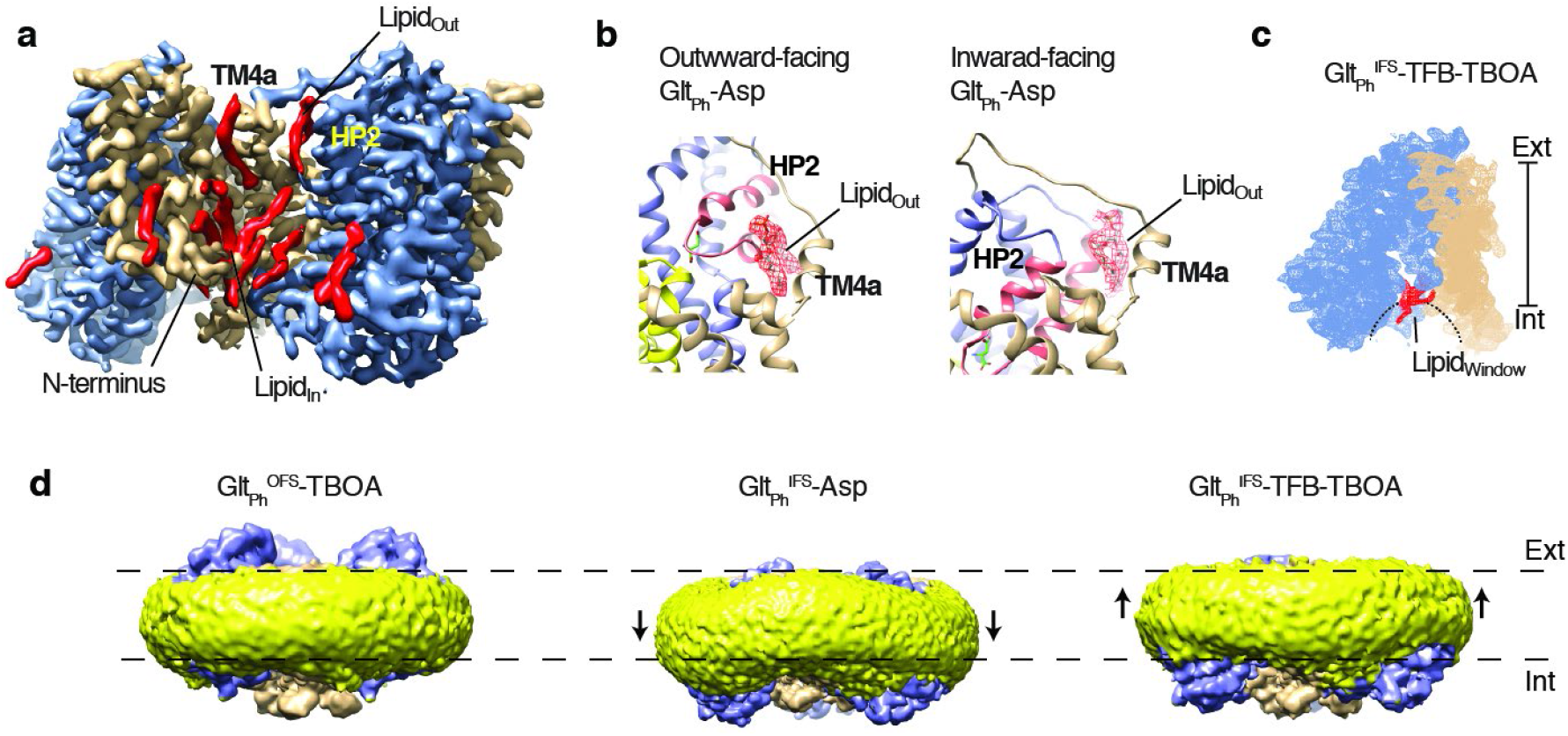
Coupling of the lipid bilayer to protein motions. **a,** Lipid densities (red) observed in protein cervices of Glt_Ph_^IFS^-Asp. Lipid molecules tucked in between the N-terminus and the rest of the scaffold (Lipid_m_) are observed in all OFS and IFS structures. **b,** lipid densities (red mesh objects, Lipidout) observed on the extracellular side of a crevice between the scaffold TM4a and HP2 in both outward- (PDB code 6UWF) and inward-facing Glt_Ph_ bound to L-asp. **c,** Density map of a Glt_Ph_^IFS^-TFB-TBOA protomer, with the lipid density in the window between the transport domain and scaffold colored red (Lipidwindow). **d**, Density map of Glt_Ph_^OFS^-TBOA, Glt_Ph_^IFS^-Asp, and Glt_Ph_^IFS^-TFB-TBOA in nanodiscs viewed in the membrane plain. Density corresponding to the transport and scaffold domains are colored blue and wheat, respectively. Density corresponding to the nanodisc is colored yellow. Black arrows mark deviations of the nanodiscs from the planar structures.

We also find lipid moieties, structured to various degrees, in the crevices between the scaffold and transport domains (**Figure 5a)**. Of these, the most notable one is inserted between the N-terminal arm of HP2 and the scaffold TM4a (LipidOut **Figure 5a, b**). Interestingly, we observe lipids at almost the same location in the outward- (Huang *et al*., 2019) and inward-facing L-asp-bound transporters (**Figure 5b**). The lipid packs similarly against TM4a in the OFS and IFS but interacts differently with HP2: near the tip and the extracellular base, respectively. It is not yet clear whether during the outward-to-inward transition, as HP2 slides past TM4a, the lipid is temporarily displaced or disordered. Interestingly HP2 opening in the OFS, as seen in Glt_Ph_^OFS^-TBOA, and the IFS, as seen in the current study, requires displacement of LipidOut. Thus, the lipid molecules at this site could modulate gating and the translocation dynamics, affecting both substrate affinity and transport rate. In Glt_Ph_^IFS^-TFB-TBOA and Glt_Ph_^IFS^-Apo-open structures, the transport domain leans away from the scaffold far enough to open a window between the two domains that connects the interior of the bilayer to the solvent-filled crevice on the cytoplasmic side of the transporter (**Figure 5c**). We observe excess densities in the opening, suggesting that lipids enter the space at a position structurally symmetric to LipidOut (LipidWindow, **Figure 5c**).

Perhaps most strikingly, we observe overall distortions of the nanodisc correlated to the position of the transport domain (**Figure 5d, Movie 3**). The nanodisc is nearly flat in the structure of the Glt_Ph_^OFS^-TBOA, in which the hydrophobic regions of the transport domain and the scaffold are aligned. In Glt_Ph_^IFS^-Asp, the transport domain is positioned at the sharpest angle to the membrane normal (**Figure 1a**), and its hydrophobic region descends the furthest toward the cytoplasm. The resulting hydrophobic mismatch between the scaffold and transport domains leads to membrane bending to accommodate both, as has been predicted by computational studies (Zhou *et al*., 2019). In other inward-facing structures, where the transport domains swing out, the hydrophobic regions are positioned further up toward the extracellular side of the membrane leading to reduced membrane bending. In the Glt_Ph_^IFS^-TFB-TBOA structure, the membrane bends in the reverse direction as the swing of the transport domains brings their hydrophobic regions above those of the scaffold (**Figure 5d, Movie 3**).

## Discussion

The series of structures that we have determined by Cryo-EM suggest that both substrate translocation and substrate gating in the IFS require movements of the transport domain through membrane bilayer. One remarkable aspect of the observed structural transitions is the conformational plasticity of HP2 and the interface between the transport domain and scaffold, which differ in each functional intermediate of the transporter. Our recent studies suggest that both translocation of the transport domain and substrate release into the cytoplasm are the slow steps of the Glt_Ph_ functional cycle (Oh e Boudker, 2018; Huysmans *et al*., 2020). Most strikingly, subtle packing mutations in HP2 at sites distant from the substrate-binding site decrease substrate affinity in the OFS and IFS, and increase the dynamics of the OFS to IFS transitions (Huysmans *et al*., 2020).

Our structures show that the release of Na^+^ ions and L-asp requires movement of the transport domain, mediated by conformational changes of HP2 and the HP2/TM8a-scaffold interface. The complexity of these conformational events may explain why the substrate binding and release are slow in the IFS (Oh e Boudker, 2018), while fast in the OFS (Hanelt *et al*., 2015). Notably, kinetic studies showed that the release (and binding) of one Na^+^ ion in the IFS, most likely Na2, is rapid (Oh e Boudker, 2018). Thus, it is likely that the release of Na2 requires little structural change, limited at most to the change observed in the Glt_Ph_^IFS^-TBOA structure. Our structural data further suggest that mutations in HP2 may increase the substrate dissociation rate in the IFS by increasing the dynamics of the hairpin and the hairpin/scaffold interface.

Single-molecules studies of the OFS to IFS translocation dynamics showed that the rate-limiting high-energy transition state most likely resembles the IFS structurally and that the transport domain might make multiple attempts to achieve a stable observable IFS (Huysmans *et al*., 2020). These studies suggest that multiple IFS conformations exist and are separated by significant energetic barriers. While our structures most likely represent the lowest-energy states, their multiplicity in itself supports the existence of a complex conformational ensemble sampled by the transporter in the IFS.

The observed functional large-scale domain movements are accompanied by significant alterations of the structure of the surrounding membranes and some of the well-structured annular lipids. In general, it appears that all indentations and crevices on the surface of the protein open to the bilayer and large enough to accommodate hydrocarbon chains are well-occupied by lipids even in the absence of specific interactions between the headgroups and protein moieties. The density for some of the lipids, such as LipidIn (**Figure 5a and Figure 5 Supplementary Figure 1**), is very well resolved, and both the location and structures of the lipid molecules are conserved in all resolved protein complexes. It is unclear, however, whether they are structurally immobilized or exchange rapidly with the surrounding bulk lipids.

Other lipids, in particular extracellular Lipid_out_, present in the OFS and IFS, and the symmetric cytoplasmic Lipid_window_ observed in the IFS, have to move in and out of their binding sites during the transport cycle. Interestingly, LipidOut is inserted between the N-terminal arm of HP2 and the scaffold in both the OFS and IFS. Thus, the HP2 arm shows a conserved behavior in the two states: When HP2 occludes the L-asp binding site, LipidOut fills the space between the arm and the scaffold, but when HP2 opens during gating, the arm interacts directly with the scaffold displacing the lipid. In contrast, LipidWindow, in the IFS, moves in when HP2 opens to release the substrate and moves out when HP2 closes. Such intimate involvement of lipids suggests that they have the potential to regulate both substrate affinity and conformational dynamics of the transporter. So far, however, only modest effects of specific lipids on Glt_Ph_ transport activity have been reported (Mcilwain *et al*., 2015). Interestingly, in mammalian EAAT1 and ASCT2, similar space between the N-terminal arm of HP2 and scaffold is observed in the OFS, and IFS (Canul-Tec *et al*., 2017; Guskov *et al*., 2018; Yu *et al*., 2019), and likely can accommodate lipids. These proteins are known to be regulated by lipids, particularly by the arachidonic acid (Zerangue *et al*., 1995; Fairman *et al*., 1998; Tzingounis *et al*., 1998), and the identified lipid-binding sites might mediate these effects.

The lipid bilayer bending to accommodate conformational change from the OFS to IFS has been observed in recent molecular dynamics simulations (Zhou *et al*., 2019). The computational study suggests that the energy penalty associated with bilayer bending might be as large as 6-7 kcal/mol protomer. Our results show that not only the OFS to IFS transitions, but also the substrate release in the IFS involve changes in membrane deformation. Thus, large energetic costs of membrane bending might accompany glutamate transporter functional cycle, suggesting that the physical properties of lipid bilayers, such as thickness and stiffness (Lundbaek *et al*., 2010; Bruno *et al*., 2013; Rusinova *et al*., 2014), might have large impacts on transporter function.

## Materials and methods

### Glt_Ph_ expression, purification and crosslinking

The fully functional seven-histidine mutant of Glt_Ph_ that has been used in previous studies and that is referred to as wildtype (WT) for brevity, and the K55C/C321A/A364C Glt_Ph_ mutant were expressed as C-terminal His8 fusions and purified as described previously (Yernool *et al*., 2004). Briefly, the plasmids were transformed into *E. coli* DH10-B cells (*Invitrogen*). Cells were grown in LB media supplemented with 0.2 mg/l of ampicillin (*Goldbio*) at 37 °C until OD_600_ of 1.0. Protein expression was induced by adding 0.2 % arabinose (*Goldbio*) for 3 hr at 37 °C. The cells were harvested by centrifugation and resuspended in 20 mM Hepes, pH 7.4, 200 mM NaCl, 1 mM L-asp, 1 mM EDTA. The suspended cells were broken using Emulsiflex C3 high pressure homogenizer (*Avestin Inc.*) in the presence of 0.5 mg/ml lysozyme (*Goldbio*) and 1 mM phenylmethanesulfonyl fluoride (PMSF, *MP Biomedicals*). After centrifugation for 15 min at 5000 g at 4 °C to remove the debris, membranes were pelleted by centrifugation at 125,000 g for 60 min. The membranes were homogenized in 20 mM Hepes, pH 7.4, 200 mM NaCl, 1 mM L-asp, 10 mM EDTA, 10 % sucrose and pelleted again by centrifugation at 125,000 g for 60 min.

The washed membranes were collected and solubilized in Buffer A, containing 20 mM Hepes, pH7.4, 200 mM NaCl, 1 mM L-asp, supplemented with 40 mM n-dodecyl-β-D-maltopyranoside (DDM, *Anatrace, Inc*.) at 8 ml per gram of membranes for 2 hours at 4 °C. The mixture was clarified by ultracentrifugation for 60 min at 125,000 g, the supernatant was incubated with Ni-NTA resin (*Qiagen*) pre-equilibrated in buffer A with gentle shaking for 2 hr at 4 °C. The resin was washed with 5 volumes of Buffer A with 1 mM DDM and 25 mM imidazole, the protein was eluted in the same buffer containing 250 mM imidazole. The eluted protein was concentrated using concentrators with 100 kDa MW cutoff (*Amicon*). The (His)8-tag was cleaved by thrombin (*Sigma*) using 20 U per 1 mg Glt_Ph_ in the presence of 5 mM CaCl2 at room temperature overnight. The reaction was stopped by addition of 10 mM EDTA and 1 mM PMSF. For the WT Glt_Ph_, the protein was further purified by size exclusion chromatography (SEC) in buffer A and 1mM DDM. The eluted protein was concentrated and used immediately for nanodisc reconstitution.

Prior to crosslinking, the K55C/C321A/A364C mutant protein was reduced with 5 mM Tris(2-carboxyethyl)phosphine (TCEP) at room temperature for 1 hr. Reduced K55C/C321A/A364C Glt_Ph_ at concentrations below 1 mg/mL was incubated with 10-fold molar excess of HgCl2 for 15 min at room temperature. The protein was concentrated to under 1 ml and purified by SEC in buffer A supplemented with 1mM DDM. The elution peak fractions were collected and concentrated. The protein concentration was determined by UV absorbance at 280 nm using extinction coefficient of 57,400 M^-1^ cm^-1^ and MW of 44.7 kDa. To check availability of free thiols after crosslinking, proteins were incubated with 5-fold molar excess of fluoroscein-5-maleimide (F5M). Fluorescent F5M-labeled proteins were imaged on SDS-PAGE under blue illumination and stained with Coomassie blue.

### Reconstitution of Glt_Ph_ into nanodiscs

Membrane scaffold protein MSP1E3 was expressed and purified from *E. coli* and Glt_Ph_ was reconstituted into lipid nanodiscs as previously described, with modifications (Ritchie *et al*., 2009). Briefly, *E. coli* polar lipid extract and egg phosphatidylcholine in chloroform (*Avanti*) were mixed at 3:1 (w:w) ratio and dried on rotary evaporator and under vacuum overnight. The dried lipid film was resuspended in buffer containing 20 mM Hepes/Tris, pH 7.4, 200 mM NaCl, 1 mM L-asp and 80 mM DDM by 10 freeze/thaw cycles resulting in 20 mM lipid stock. The purified Glt_Ph_ protein in DDM was mixed with MSP1E3 and lipid stock at 0.75:1:50 molar ratio at the final lipid concentration of 5 mM and incubated at 21 °C for 30 min. Biobeads SM2 (*Bio-Rad*) were added to one third of the reaction volume and the mixture was incubated at 21 °C for 2 hr on a rotator. Biobeads were replaced and incubated at 4 °C overnight. The sample containing Glt_Ph_^IFS^ reconstituted into the nanodiscs in the presence of 1 mM L-asp was cleared by centrifugation at 100,000 g and purified by SEC using a Superose 6 Increase 10/300 GL column (GE Lifesciences) pre-equilibrated with buffer containing 20 mM Hepes/Tris, pH 7.4, 200 mM NaCl and 1 mM L-asp. The peak fractions corresponding to Glt_Ph_^IFS^-containing nanodiscs were collected for Cryo-EM imaging. To prepare substrate-free WT Glt_Ph_ and Glt_Ph_^IFS^ in nanodiscs, the reconstitution mixtures were cleared by centrifugation at 100,000 g, diluted with 10 x volume of buffer containing 20 mM Hepes/Tris, pH 7.4, and 50 mM choline chloride, and concentrated using 100 kDa cutoff concentrator. After repeating the procedure twice, substrate-free transporters in nanodiscs were purified by SEC in the same buffer. The peak fractions were collected and immediately supplemented with buffers containing 200 mM NaCl and 10 mM DL-TBOA, 200 mM NaCl and 10 mM TFB-TBOA, or 200 mM NaCl. The presence of the MSP1E3 and Glt_Ph_ proteins in the samples was confirmed by SDS-PAGE. Negative staining electron microscopy was used to confirm the formation and the homogeneity of the nanodisc samples.

### Cryo-EM data collection

To prepare cryo-grids, 3.5 μL of Glt_Ph_-containing nanodiscs (7 mg/mL) supplemented with 1.5 mM fluorinated Fos-Choline-8 (*Anatrace*) were applied to a glow-discharged UltrAuFoil R1.2/1.3 300-mesh gold grid (*Quantifoil*) and incubated for 20 s under 100 % humidity at 15 °C. Grids were blotted for 2 s and plunge frozen in liquid ethane using Vitrobot Mark IV (*Thermo Fisher Scientific*). For the WT Glt_Ph_ in the presence of DL-TBOA (Glt_Ph_^OFS^-TBOA), Glt_Ph_^IFS^ in the presence of TFB-TBOA (Glt_Ph_^IFS^-TFB-TBOA), and Glt_Ph_^IFS^ in the presence of 200 mM Na^+^ ions only (Glt_Ph_^IFS^-NaCl), the Cryo-EM imaging data were acquired using a Titan Krios microscope (*Thermo Fisher Scientific*) at New York Structural Biology Center operated at 300 kV with a K2 Summit detector with a calibrated pixel size of 1.07325 Å/pixel. A total dose of 68.55 e^-^/Å^2^ (Glt_Ph_^OFS^-TBOA, Glt_Ph_^IFS^-NaCl), or 68.70 e^-^/Å^2^ (Glt_Ph_^IFS^-TFB-TBOA) distributed over 45 frames (1.52 e / Å^2^/frame) was used with an exposure time of 9 s (200 ms/frame) and a defocus range of −1.5 μm to −2.5 μm. For Glt_Ph_^IFS^ in the presence of DL-TBOA (Glt_Ph_^IFS^-TBOA), Cryo-EM imaging data were acquired on a Titan Krios microscope at New York Structural Biology Center operated at 300 kV with a K2 Summit detector with a calibrated pixel size of 1.07325 Å/pixel. A total dose of 69.70 e^-^/Å^2^ distributed over 50 frames (1.52 e^-^/ Å^2^/frame) was used with an exposure time of 10 s (200 ms/frame) and a defocus range of −1.5 μm to −2.5 μm. For the Glt_Ph_^IFS^ in the presence of L-asp (Glt_Ph_^IFS^-Asp), micrographs were acquired o n a Titan Krios microscope at New York Structural Biology Center operated at 300 kV with a K2 Summit detector, using a slid width of 20 eV on a GIF Quantum energy filter with a calibrated pixel size of 1.0605 Å/pixel. A total dose of 69.30 e^-^/Å^2^ distributed over 45 frames (1.54 e^-^/ Å^2^/frame) was used with an exposure time of 9 s (200 ms/frame) and defocus range of −1.5 μm to −2.5 μm. For all samples, automated data collection was carried out using Leginon (Suloway et al., 2005).

### Image processing

The frame stacks were motion corrected using MotionCorr2 (Zheng *et al*., 2017) and contrast transfer function (CTF) estimation was performed using CTFFIND4 (Rohou e Grigori eff, 2015). All further processing steps were done using RELION 3.0 unless otherwise indicated (Zivanov *et al*., 2018). Dogpicker (Voss *et al*., 2009) as part of the Appion processing package (Lander *et al*., 2009) was used for reference-free particle picking. Picked particles were then extracted and subjected to 2D classification to generate 2D class-averages which were used as templates for automated particle picking in Relion. The particles were extracted using a box size of 275 Å with 2x binning and subjected to 2 rounds of 2D classification ignoring CTFs until the first peak.

For Glt_Ph_^IFS^-Asp, Glt_Ph_^IFS^-TBOA, Glt_Ph_^IFS^-TFB-TBOA, and for the Glt_Ph_^OFS^-TBOA, particles selected from 2D classification were re-extracted without binning and further classified into 6 classes without enforcing symmetry using initial models generated in

CryoSPARC (Punjani et al., 2017) and filtered to 40 Å. Particles from the best classes showing trimeric transporter arrangements were subjected to 3D refinement applying C3 symmetry. After conversion, the refinement was continued with a mask excluding the nanodisc. To further improve the resolution of the maps, the particles after 3D refinement were subject to an additional round of 3D classification without alignment with C3 symmetry and T=4 applying a mask to exclude the nanodisc. Particles from the best class were subjected to further masked refinement and CTF refinement. A masked refinement following CTF refinement yielded final maps with the following resolution: 3.05 Å (Glt_Ph_^IFS^-Asp), 3.71 Å (Glt_Ph_^IFS^-TFB-TBOA), 3.66 Å (Glt_Ph_^IFS^-TBOA), 3.66 Å (Glt_Ph_ - TBOA). The resolution limits of the refined maps were assessed using Relion postprocessing and gold standard FSC value 0.143 using masks that excluded the nanodiscs. To test for potential conformational heterogeneity, we re-processed the datasets with no symmetry applied (C1). The obtained maps showed slightly lower resolution but no detectable difference when compared to the results from the C3 refinement. We also processed the datasets with symmetry expansion (C3) and did not find additional conformations.

During processing of the data for Glt_Ph_^IFS^-NaCl, 529,155 particles selected from 2D classification were re-extracted without binning and were subjected to 3D classification with K=1 and no symmetry applied, using Glt_Ph_^IFS^-Asp map as the initial model. The same particles were subject to 3D refinement with C3 symmetry. After conversion, the refinement was continued with a mask to exclude the nanodisc, resulting in a 3.56 Å resolution map. To probe for conformational heterogeneity, we performed symmetry expansion implemented in Relion (Scheres, 2016). 1,587,465 protein subunits were rotated to the same position and subjected to a focused 3D classification without alignment with T=40 into 10 classes. The local mask was generated using Chain A of PDB model 3KBC and included only densities from one subunit of the refence map. Two different conformations were observed. From the 10 classes, 5 classes showed a conformation identified as Glt_Ph_^IFS^-Na and 5 classes showed a different conformation identified as Glt_Ph_^IFS^-Apo-open. The best Glt_Ph_^IFS^-Na^+^ class (191,349 particles), which contained 12 % of the symmetry expanded protomers and the best Glt_Ph_^IFS^-Apo-open class (148,582 particles), which contained 9 % of the symmetry expanded particles, were separately subjected to a final focused 3D refinement with C1 using a mask to exclude the nanodisc. The local angular searches in this refinement were conducted only around the expanded set of orientations to prevent contributions from the neighbor subunits in the same particle. The resulting maps were postprocessed in Relion using the same mask as in 3D classification after symmetry expansion. The final resolution at gold standard FSC value 0.143 was estimated as 3.52 Å for the Glt_Ph_^IFS^-Apo-open map and 3.66 Å for Glt_Ph_^IFS^-Na map. Local resolution variations were estimated using ResMap (Kucukelbir *et al*., 2014). After symmetry expansion with C3, we also tried to first subtract the density outside of one Glt_Ph_ subunit and then perform 3D classification without alignment on the subtracted particles. The signal subtraction did not further improve the 3D classification and the 3D refinement.

### Model building and refinement

For atomic model building from Glt_Ph_^IFS^-Asp, Glt_Ph_^IFS^-TBOA, and Glt_Ph_^IFS^-TFB-TBOA maps, crystal structure of Glt_Ph_ in the IFS (accession code 3KBC) was docked into the density maps using UCSF Chimera (Pettersen *et al*., 2004). For the WT Glt_Ph_^OFS^-TBOA, crystal structure of Glt_Ph_ in the OFS (accession code 2NWW) was docked into the density. For Glt_Ph_^IFS^-Na or Glt_Ph_^IFS^-Apo-open, one subunit of 3KBC was docked into the density. After the first rounds of the real-space refinement using Phenix (Afonine *et al*., 2010), miss-aligned regions were manually rebuilt and missing side chains and residues were added in COOT (Emsley *et al*., 2010). 1-palmitoyl-2-oleoyl-sn-glycero-3-phosphoethanolamine (POPE) was used as a model lipid and placed into the excess densities which resembled lipid molecules. The acyl chains or ethanolamine heads were truncated to fit the visible densities. Models were iteratively refined applying secondary structure restraints and validated using Molprobity (Chen *et al*., 2010). For further cross validation and to check for overfitting, all atoms of each model were randomly displaced by 0.3 Å and each resulting model was refined against the first half-map obtained from processing. FSC between the refined models and the half-maps used during the refinement were calculated and compared to the FSC between the refined models and the other halfmaps. In addition, the FSC between the refined model and sum of both half-maps was calculated. The resulting FSC curves were similar showing no evidence of overfitting.

## SUPPLEMENTARY INFORMATION

**Figure 1 Supplementary Figure 1.**
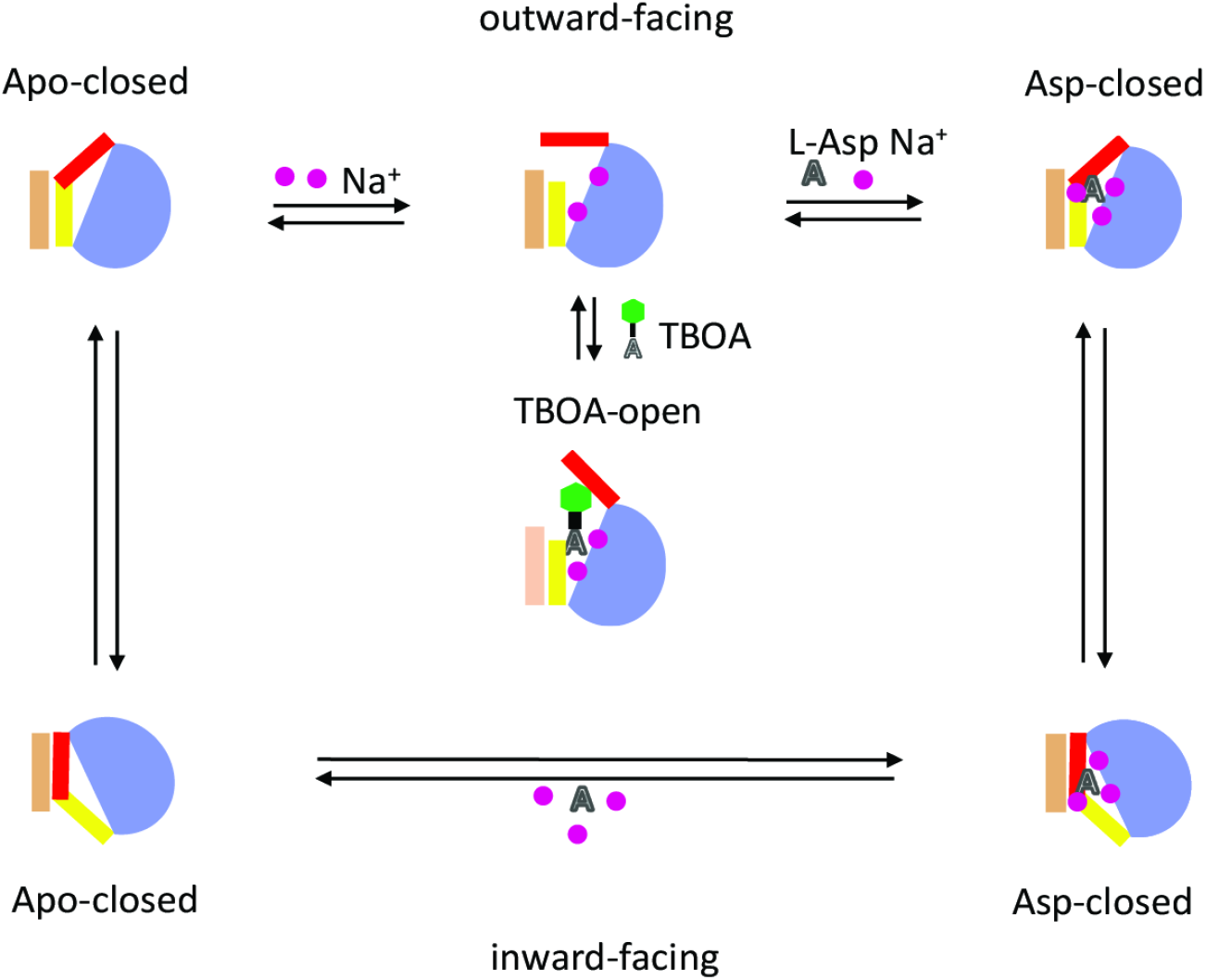
Schematic representation of the elevator mechanism of transport by Glt_Ph_. The scaffold domain is in wheat, and the transport domain is in gray. HP1 and HP2 are yellow and red, respectively. Substrate L-asp is represented as a letter A and the benzyl group of the blocker TBOA is shown as a green hexagon. Three symported Na^+^ ions are shown as purple circles.

**Figure 1 Supplementary Figure 2.**
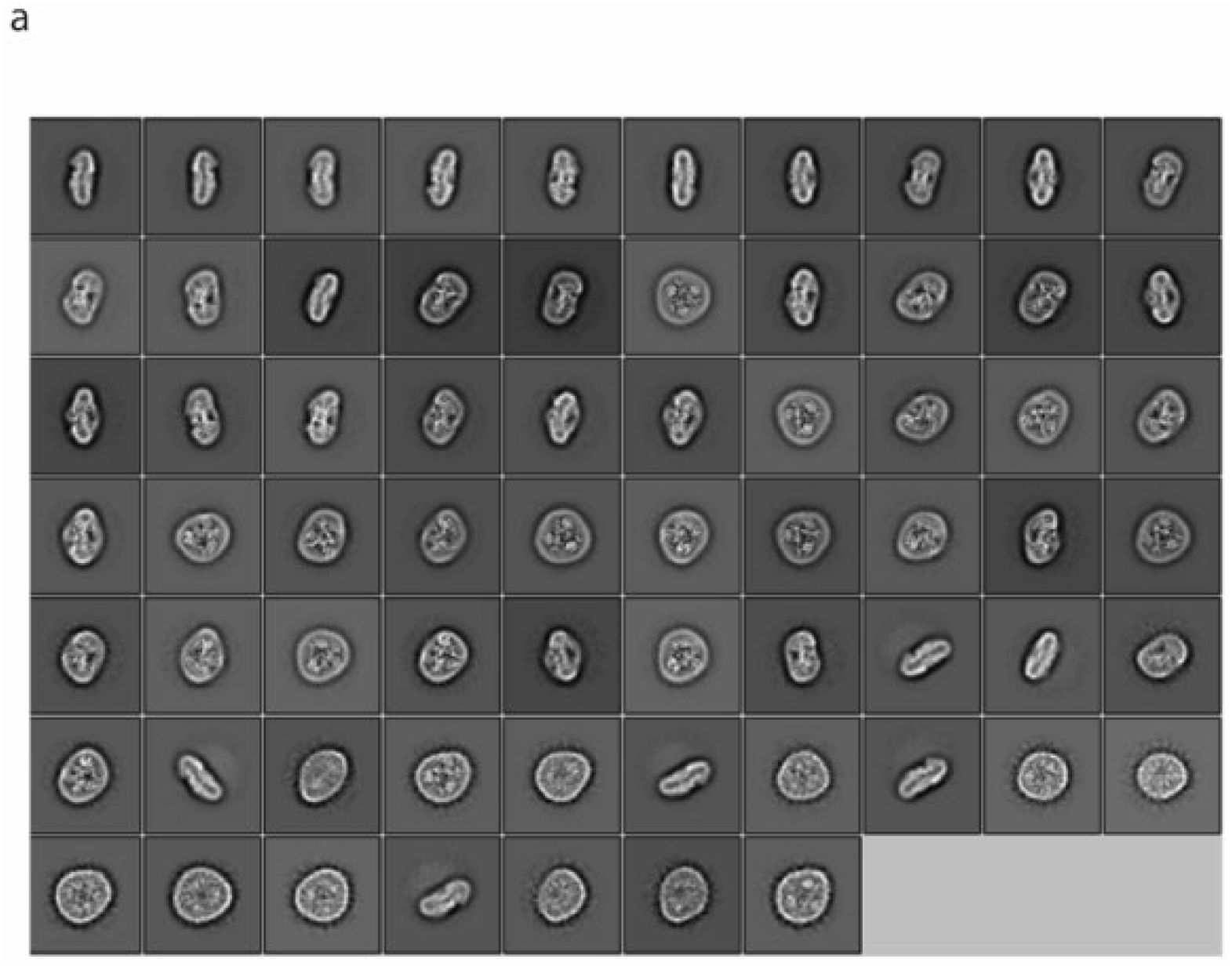
Cryo-EM data processing. An example of selected 2D classes depicting the trimeric Glt_Ph_^IFS^-Asp transporter in lipid nanodiscs. Box size is 275 Å

**Figure 1 Supplementary Figure 3.**
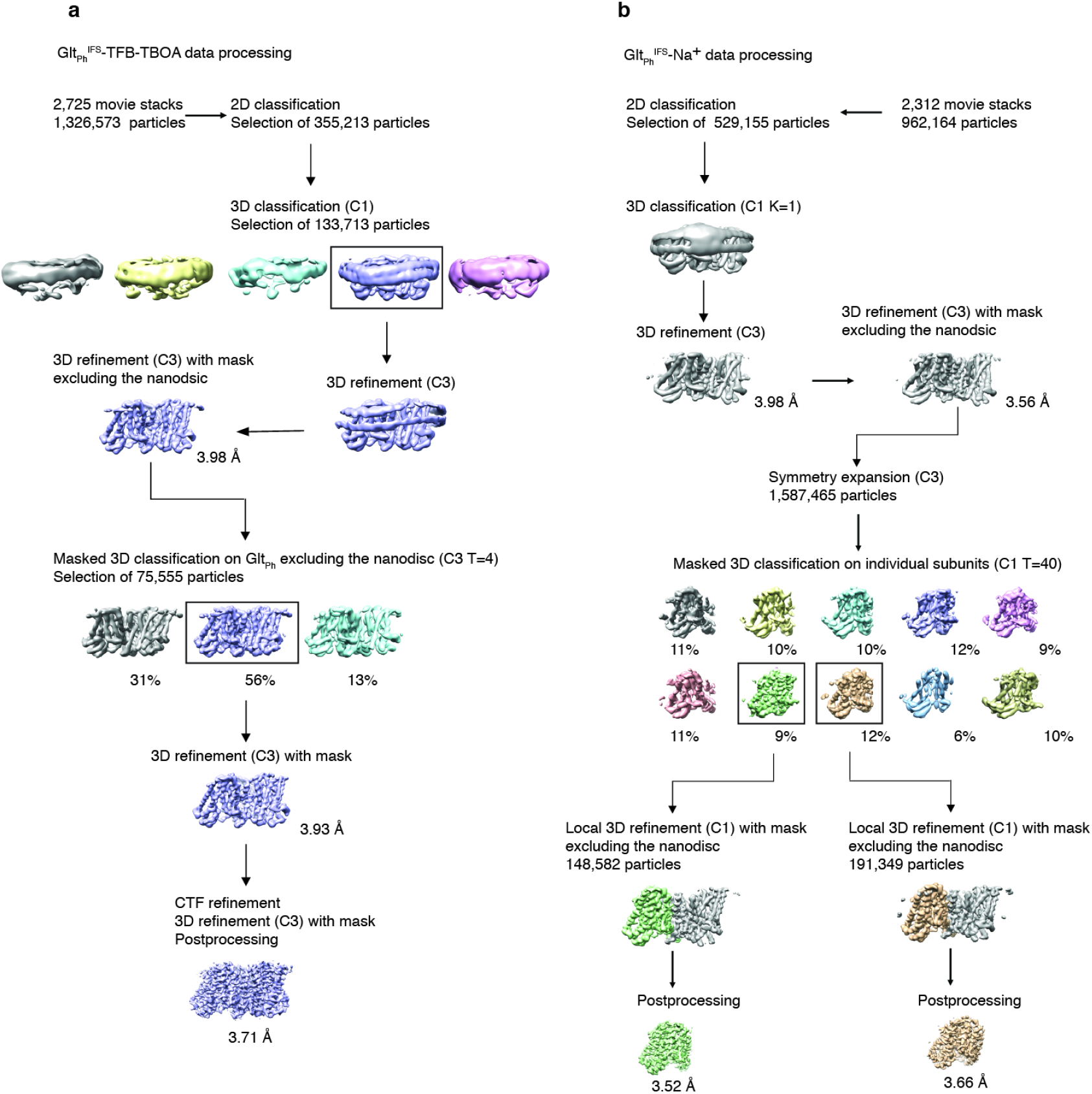
Data processing flowchart for Glt_Ph_^IFS^-TFB-TBOA (a) and Glt_Ph_^IFS^-Na, and Glt_Ph_^IFS^-Apo-open (b). Data processing for Glt_Ph_^IFS^-Asp, Glt_Ph_^IFS^ -TBOA, and Glt_Ph_^OFS^-TBOA followed the same scheme as Glt_Ph_^IFS^-TFB-TBOA.

**Figure 1 Supplementary Figure 4.**
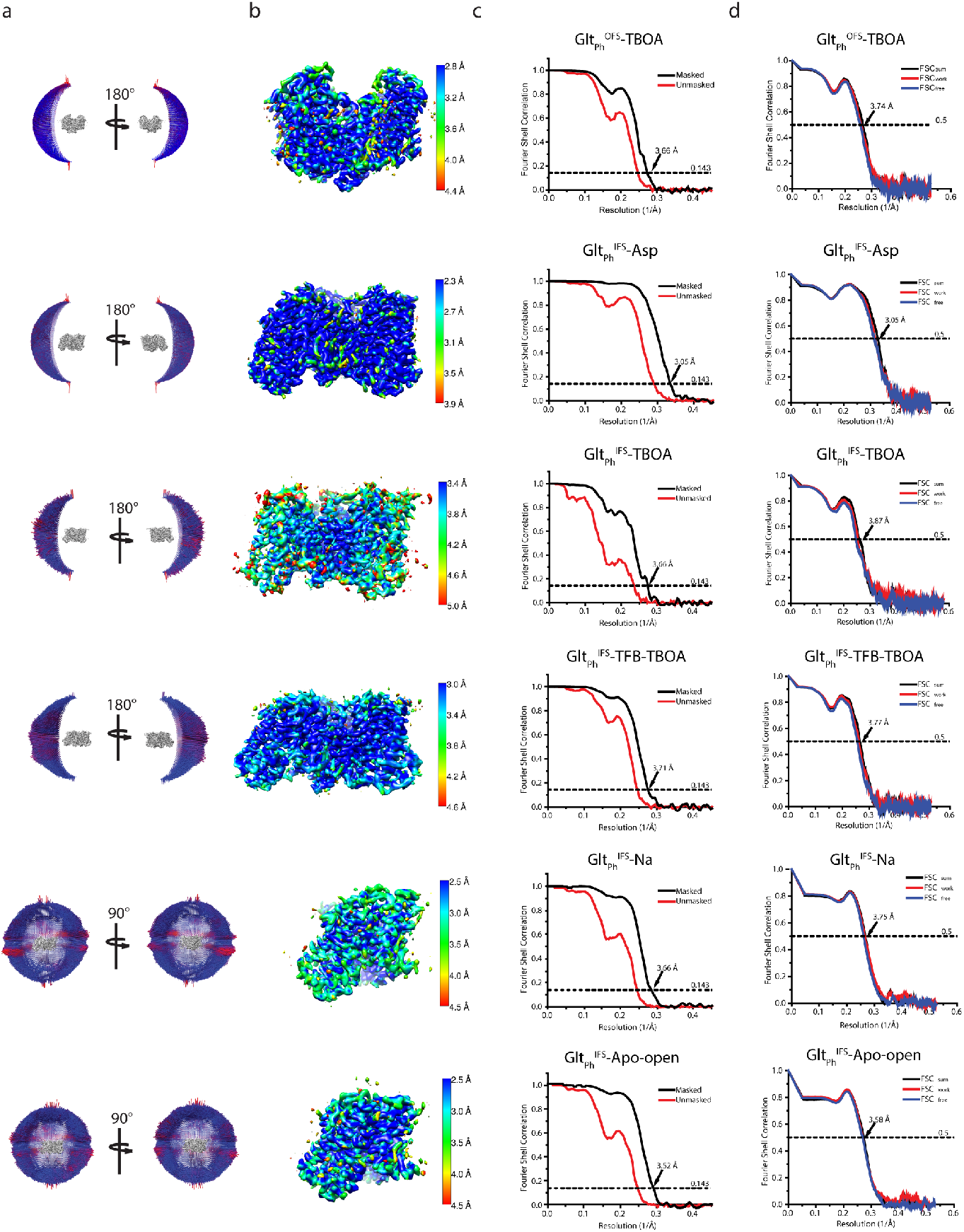
Cryo-EM imaging and data processing validation. **a**, Angular distribution of particles contributing to the final reconstitutions. The number of views at each angular orientation is represented by the length and color of cylinders, where red indicates more views. **b**, Final maps after Relion post-processing colored according to the local resolution estimation using ResMap. **c**, Fourier shell correlation (FSC) curves indicating the resolution at the 0.143 thresholds of the final masked (black) and unmasked (orange) maps. **d**, FSC curves from cross-validation of the refined models compared to the masked half-map 1 (orange traces: FSCwork, used during validation refinement), masked half map 2 (blue traces: FSCfree, not used during validation refinement), and the masked summed map (black traces: FSCsum).

**Figure 2 Supplementary Figure 1.**
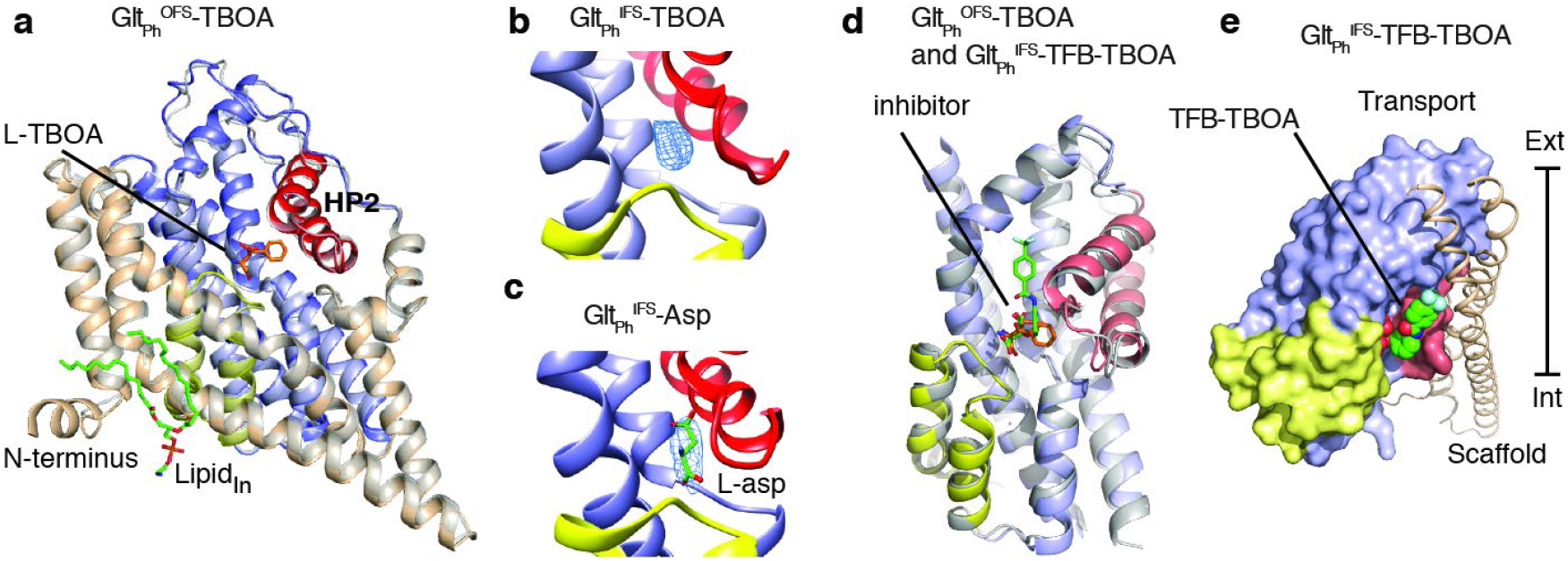
**a.** Superimposition of WT Glt_Ph_^OFS^-TBOA Cryo-EM (colored) and crystal structures (grey, accession code 2NWW). L-TBOA modeled in the crystal structure (orange), and a lipid molecule modeled in the Cryo-EM structure (green) are shown as sticks. The rest of the protein is colored as in Figure 1. **b, c,** Close-up views of the substrate-binding site in Glt_Ph_^IFS^ -TBOA (**b**) and Glt_Ph_^IFS^ -Asp (**c**). Excess densities are shown as blue mesh objects. The modeled L-asp molecule is shown in stick representation. **d,** Superimposition of the transport domains of Glt_Ph_^IFS^-TFB-TBOA (colors) and WT Glt_Ph_^OFS^-TBOA (grey). Bound L-TBOA and TFB-TBOA are shown in stick representation and colored orange and green, respectively. **e**, Glt_Ph_^IFS^-TFB-TBOA protomer viewed in membrane plain with transport and scaffold domains shown in surface and cartoon representations, respectively. TFB-TBOA (green spheres) protrudes from the binding pocket toward the domain interface.

**Figure 3 Supplementary Figure 1.**
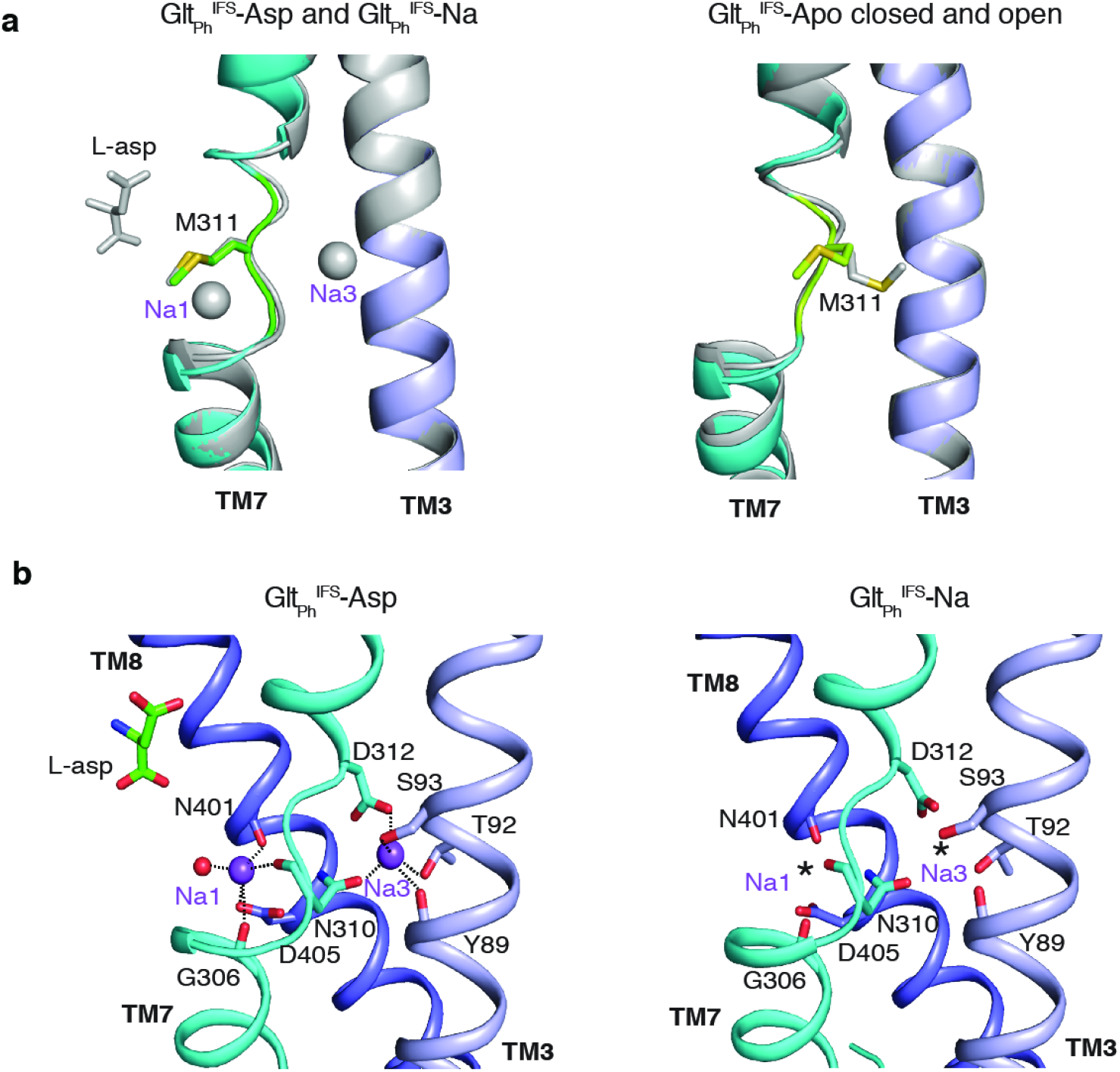
Na^+^-binding sites in the Glt_Ph_^IFS^ in Apo and Na^+^-bound states. **a**, The superimposition of Glt_Ph_^IFS^-Na and Glt_Ph_^IFS^-Asp (left) and Glt_Ph_^IFS^-Apo-open and Apo-closed (right) show similar conformations of NMD motif, involved in coordinating Na^+^ ions at Na1 and Na3 sites. **b**, Structures of Na1 and Na3 sites in Glt_Ph_^IFS^-Asp and Glt_Ph_^IFS^-Na. Coordinating moieties within 3 Å of the ions are emphasized by dotted lines in Glt_Ph_^IFS^-Asp. Stars in Glt_Ph_^IFS^-Na represent the potentially bound Na^+^ ions, placed as in Glt_Ph_^IFS^-Asp.

**Figure 3 Supplementary Figure 2.**
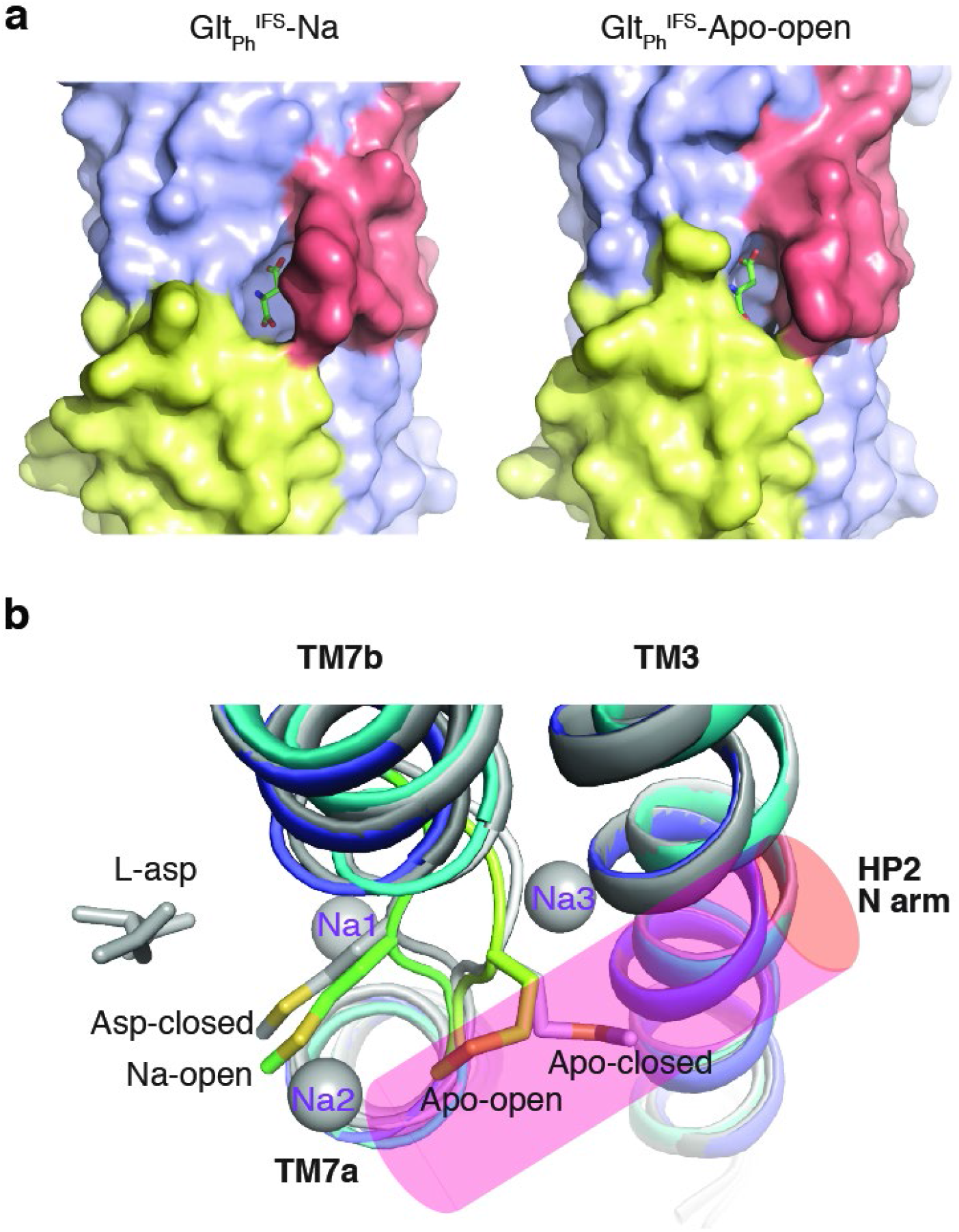
Molecular mechanism of HP2 opening and closing. **a**, Surface representation of the two open structures, Glt_Ph_^IFS^-Na and Glt_Ph_^IFS^-Apo-open with HP1 and HP2 colored yellow and red, respectively. To emphasize solvent accessibility of the substrate-binding site, L-asp is shown in stick representation at the position found in the Glt_Ph_^IFS^-Asp structure. **b,** Top view of superimposed transport domains of Glt_Ph_^IFS^-Asp (dark grey), Glt_Ph_^IFS^-Na (dark colors), Glt_Ph_^IFS^-Apo-open (light colors), and Glt_Ph_^IFS^-Apo closed (light grey). Only TM7 and TM3 are shown for clarity. Sidechains of M311 are depicted as sticks.

**Figure 4 Supplementary Figure 1.**
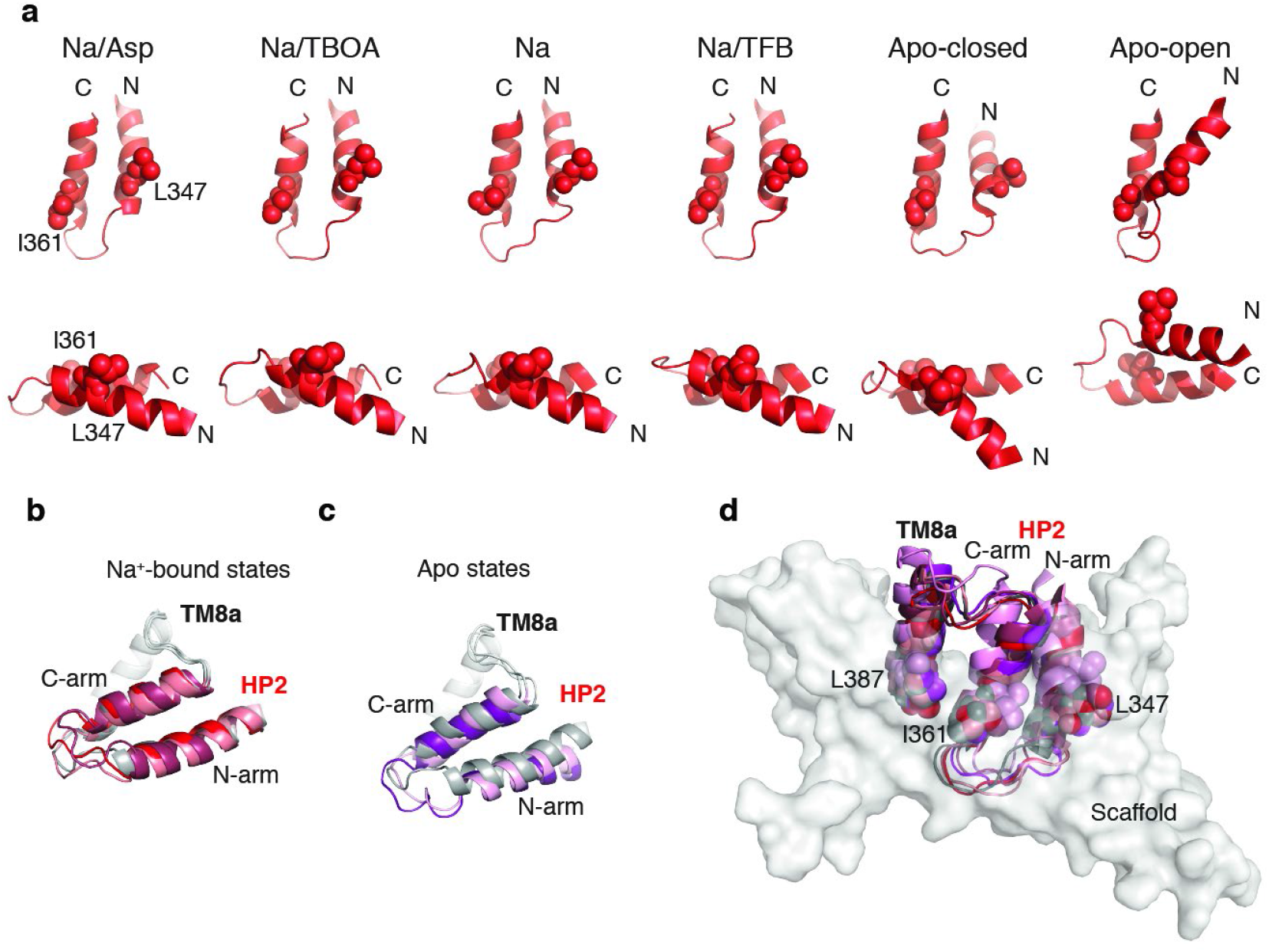
Structural plasticity of HP2 and of the inter-domain interface. **a and b,** Cartoon representation of HP2/TM8a helices aligned on TM8a for structures with (**a**) and without (**b**) bound Na1 and Na3. Glt_Ph_^IFS^-Asp (grey) is shown in both panels for reference. Glt_Ph_^IFS^-TBOA is colored salmon, Glt_Ph_^IFS^-Na red, Glt_Ph_^IFS^-TFB-TBOA berry, Glt_Ph_^IFS^-Apo-closed (pink), Glt_Ph_^IFS^-Apo-open (purple).

**Figure 5 Supplementary Figure 1.**
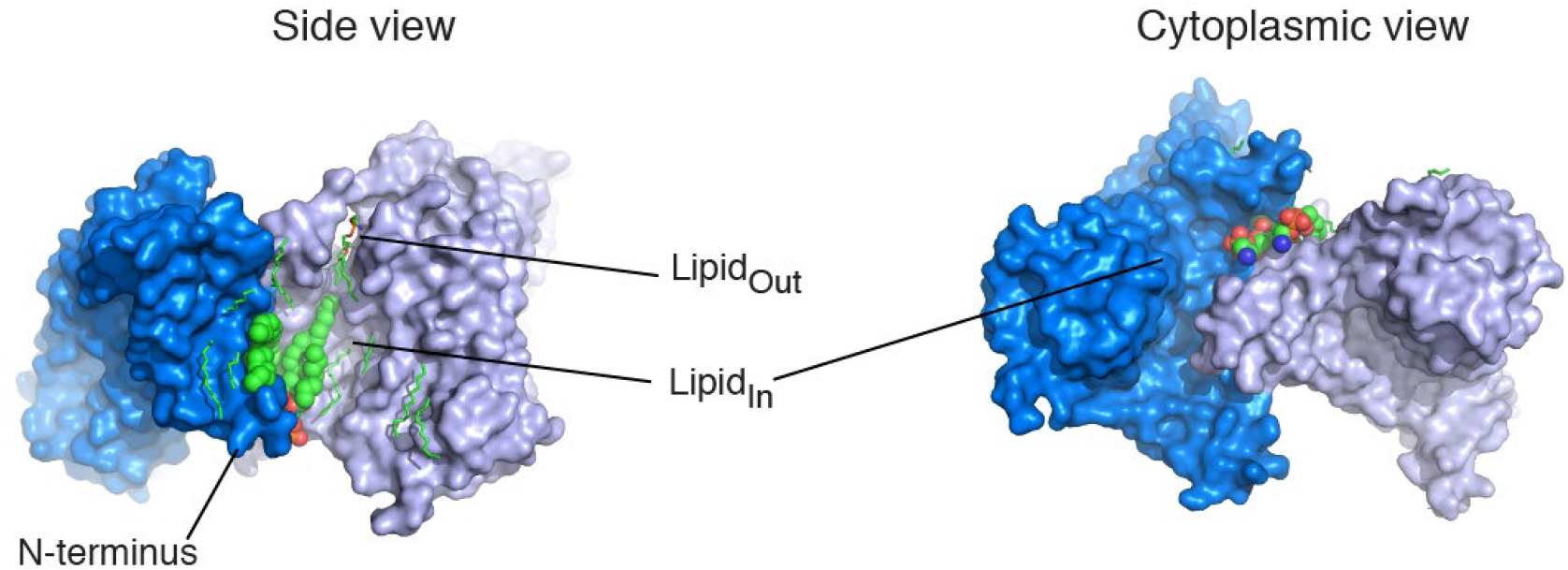
Structured lipids. Two Glt_Ph_^IFS^-Asp protomers are shown in surface representation and colored marine, and light blue and viewed in membrane plain (left), or from the cytoplasmic side (right). Structured lipids are colored by atom type and shown as sticks, except the two structured lipid molecules at the N-terminus (LipidIn), which are shown as spheres.

### ACKNOWLEDGEMENTS

The authors thank members of the Boudker lab for helpful discussions. Drs Biao Qiu, Maria E Falzone, and Jan Rheinberger for helpful suggestions on cryo-EM data processing, and R. Lea Sanford for insightful discussions. This work was supported by NIH Grants R01NS064357 and R37NS085318 (to OB). All EM data collections were carried out at the Simons Electron Microscopy Center and National Resource for Automated Molecular Microscopy located at the New York Structural Biology Center, supported by grants from the Simons Foundation (349247), NYSTAR, and the NIH National Institute of General Medical Sciences (GM103310) with additional support from Agouron Institute (F00316) and NIH S10 OD019994-01. Initial negative stain screening was performed at the Weill Cornell Microscopy and Image Analysis Core Facility, with the help of Dr. L Cohen-Gould.

## REFERENCES

Afonine, P. V. et al. phenix.model_vs_data: a high-level tool for the calculation of crystallographic model and data statistics. J Appl Crystallogr, v. 43, n. Pt 4, p. 669–676, Aug 1 2010. ISSN 0021-8898 (Print) 0021-8898 (Linking). Disponível em: < https://www.ncbi.nlm.nih.gov/pubmed/20648263 >.

Akyuz, N. et al. Transport dynamics in a glutamate transporter homologue. Nature, v. 502, n. 7469, p. 114-+, Oct 3 2013. ISSN 0028-0836. Disponível em: < <Go to ISI>://WOS:000325106000042 >.

Akyuz, N. et al. Transport domain unlocking sets the uptake rate of an aspartate transporter. Nature, v. 518, n. 7537, p. 68-+, Feb 5 2015. ISSN 0028-0836. Disponível em: < <Go to ISI>://WOS:000349098000032 >.

Arkhipova, V.; Guskov, A.; Slotboom, D. J. Structural ensemble of a glutamate transporter homologue in lipid nanodisc environment. Nature Communications, v. 11, n. 1, Feb 21 2020. ISSN 2041-1723. Disponível em: < <Go to ISI>://WOS:000517991000004 >.

Boudker, O. et al. Coupling substrate and ion binding to extracellular gate of a sodium-dependent aspartate transporter. Nature, v. 445, n. 7126, p. 387–393, Jan 25 2007. ISSN 0028-0836. Disponível em: < <Go to ISI>://WOS:000243689500030 >.

Bruno, M. J. et al. Interactions of drugs and amphiphiles with membranes: modulation of lipid bilayer elastic properties by changes in acyl chain unsaturation and protonation. Faraday Discuss, v. 161, p. 461–80; discussion 563-89, 2013. ISSN 1359-6640 (Print) 1359-6640 (Linking). Disponível em: < https://www.ncbi.nlm.nih.gov/pubmed/23805753 >.

Canul-Tec, J. C. et al. Structure and allosteric inhibition of excitatory amino acid transporter 1. Nature, v. 544, n. 7651, p. 446-+, Apr 27 2017. ISSN 0028-0836. Disponível em: < <Go to ISI>://WOS:000400051900036 >.

Chen, V. B. et al. MolProbity: all-atom structure validation for macromolecular crystallography. Acta Crystallogr D Biol Crystallogr, v. 66, n. Pt 1, p. 12–21, Jan 2010. ISSN 1399-0047 (Electronic) 0907-4449 (Linking). Disponível em: < https://www.ncbi.nlm.nih.gov/pubmed/20057044 >.

Crisman, T. J. et al. Inward-facing conformation of glutamate transporters as revealed by their inverted-topology structural repeats. Proc Natl Acad Sci U S A, v. 106, n. 49, p. 20752–7, Dec 8 2009. ISSN 1091-6490 (Electronic) 0027-8424 (Linking). Disponível em: < https://www.ncbi.nlm.nih.gov/pubmed/19926849 >.

Danbolt, N. C. Glutamate uptake. Progress in Neurobiology, v. 65, n. 1, p. 1–105, Sep 2001. ISSN 0301-0082. Disponível em: < <Go to ISI>://WOS:000169390500001 >.

DechancIE, J.; Shrivastava, I. H.; Bahar, I. The mechanism of substrate release by the aspartate transporter GltPh: insights from simulations. Mol Biosyst, v. 7, n. 3, p. 832–42, Mar 2011. ISSN 1742-2051 (Electronic) 1742-2051 (Linking). Disponível em: < https://www.ncbi.nlm.nih.gov/pubmed/21161089 >.

Emsley, P. et al. Features and development of Coot. Acta Crystallogr D Biol Crystallogr, v. 66, n. Pt 4, p. 486–501, Apr 2010. ISSN 1399-0047 (Electronic) 0907-4449 (Linking). Disponível em: < https://www.ncbi.nlm.nih.gov/pubmed/20383002 >.

Erkens, G. B. et al. Unsynchronised subunit motion in single trimeric sodium-coupled aspartate transporters. Nature, v. 502, n. 7469, p. 119-+, Oct 3 2013. ISSN 0028-0836. Disponível em: < <Go to ISI>://WOS:000325106000043 >.

Fairman, W. A. et al. Arachidonic acid elicits a substrate-gated proton current associated with the glutamate transporter EAAT4. Nat Neurosci, v. 1, n. 2, p. 105–13, Jun 1998. ISSN 1097-6256 (Print) 1097-6256 (Linking). Disponível em: < https://www.ncbi.nlm.nih.gov/pubmed/10195124 >.

Focke, P. J.; Moenne-Loccoz, P.; Larsson, H. P. Opposite Movement of the External Gate of a Glutamate Transporter Homolog upon Binding Cotransported Sodium Compared with Substrate. Journal of Neuroscience, v. 31, n. 16, p. 6255–6262, Apr 20 2011. ISSN 0270-6474. Disponível em: < <Go to ISI>://WOS:000289769400037 >.

Garaeva, A. A. et al. A one-gate elevator mechanism for the human neutral amino acid transporter ASCT2. Nature Communications, v. 10, Jul 31 2019. ISSN 2041-1723. Disponível em: < <Go to ISI>://WOS:000477952600012 >.

Garaeva, A. A. et al. Cryo-EM structure of the human neutral amino acid transporter ASCT2. Nature Structural & Molecular Biology, v. 25, n. 6, p. 515-+, Jun 2018. ISSN 1545-9993. Disponível em: < <Go to ISI>://WOS:000434470800012 >.

Guskov, A. et al. The stucture of human ASCT2 neutral amino acid transporter. Acta Crystallographica a-Foundation and Advances, v. 74, p. E223–E223, Aug 2018. ISSN 2053-2733. Disponível em: < <Go to ISI>://WOS:000474406600348 >.

Hanelt, I. et al. Low Affinity and Slow Na+ Binding Precedes High Affinity Aspartate Binding in the Secondary-active Transporter GltPh. J Biol Chem, v. 290, n. 26, p. 15962-72, Jun 26 2015. ISSN 1083-351X (Electronic) 0021-9258 (Linking). Disponível em: < https://www.ncbi.nlm.nih.gov/pubmed/25922069 >.

Huang, Y. et al. Monitoring Dynamics of Large Membrane Proteins by ^19^F Paramagnetic Longitudinal Relaxation: Domain Movement in a Glutamate Transporter Homolog. bioRxiv, p. 832121, 2019. Disponível em: < https://www.biorxiv.org/content/biorxiv/early/2019/11/05/832121.full.pdf >.

Huysmans, G. H. M. et al. The high-energy transition state of a membrane transporter. bioRxiv, p. 2020.04.17.047373, 2020. Disponível em: < https://www.biorxiv.org/content/biorxiv/early/2020/04/18/2020.04.17.047373.full.pdf >.

Jensen, S. et al. Crystal structure of a substrate-free aspartate transporter. Nature Structural & Molecular Biology, v. 20, n. 10, p. 1224-+, Oct 2013. ISSN 1545-9993. Disponível em: < <Go to ISI>://WOS:000325369500015 >.

Kucukelbir, A.; Sigworth, F. J.; Tagare, H. D. Quantifying the local resolution of cryo-EM density maps. Nat Methods, v. 11, n. 1, p. 63–5, Jan 2014. ISSN 1548-7105 (Electronic) 1548-7091 (Linking). Disponível em: < https://www.ncbi.nlm.nih.gov/pubmed/24213166 >.

Lander, G. C. et al. Appion: an integrated, database-driven pipeline to facilitate EM image processing. J Struct Biol, v. 166, n. 1, p. 95–102, Apr 2009. ISSN 1095-8657 (Electronic) 1047-8477 (Linking). Disponível em: < https://www.ncbi.nlm.nih.gov/pubmed/19263523 >.

LundbaeK, J. A.; Koeppe, R. E., 2ND; Andersen, O. S. Amphiphile regulation of ion channel function by changes in the bilayer spring constant. Proc Natl Acad Sci U S A, v. 107, n. 35, p. 15427–30, Aug 31 2010. ISSN 1091-6490 (Electronic) 0027-8424 (Linking). Disponível em: < https://www.ncbi.nlm.nih.gov/pubmed/20713738 >.

McilwaiN, B. C.; VandenbERG, R. J.; Ryan, R. M. Transport rates of a glutamate transporter homologue are influenced by the lipid bilayer. J Biol Chem, v. 290, n. 15, p. 9780–8, Apr 10 2015. ISSN 1083-351X (Electronic) 0021-9258 (Linking). Disponível em: < https://www.ncbi.nlm.nih.gov/pubmed/25713135 >.

Characterization of the Inward- and Outward-Facing Substrate Binding Sites of the Prokaryotic Aspartate Transporter, GltPh. Biochemistry, v. 55, n. 49, p. 6801–6810, Dec 13 2016. ISSN 1520-4995 (Electronic) 0006-2960 (Linking). Disponível em: < https://www.ncbi.nlm.nih.gov/pubmed/27951659 >.

Oh, S.; Boudker, O. Kinetic mechanism of coupled binding in sodium-aspartate symporter GltPh. Elife, v. 7, Sep 26 2018. ISSN 2050-084x. Disponível em: < <Go to ISI>://WOS:000446601600001 >.

Pettersen, E. F. et al. UCSF Chimera--a visualization system for exploratory research and analysis. J Comput Chem, v. 25, n. 13, p. 1605–12, Oct 2004. ISSN 0192-8651 (Print) 0192-8651 (Linking). Disponível em: < https://www.ncbi.nlm.nih.gov/pubmed/15264254 >.

Reyes, N.; Ginter, C.; Boudker, O. Transport mechanism of a bacterial homologue of glutamate transporters. Nature, v. 462, n. 7275, p. 880–885, Dec 17 2009. ISSN 0028-0836. Disponível em: < <Go to ISI>://WOS:000272795400034 >.

Reyes, N.; Oh, S.; Boudker, O. Binding thermodynamics of a glutamate transporter homolog. Nature Structural & Molecular Biology, v. 20, n. 5, p. 634-+, May 2013. ISSN 1545-9993. Disponível em: < <Go to ISI>://WOS:000318617000018 >.

RiedereR, E. A.; Valiyaveetil, F. I. Investigation of the allosteric coupling mechanism in a glutamate transporter homolog via unnatural amino acid mutagenesis. Proceedings of the National Academy of Sciences of the United States of America, v. 116, n. 32, p. 15939–15946, Aug 6 2019. ISSN 0027-8424. Disponível em: < <Go to ISI>://WOS:000478971900034 >.

Ritchie, T. K. et al. Chapter 11 - Reconstitution of membrane proteins in phospholipid bilayer nanodiscs. Methods Enzymol, v. 464, p. 211–31, 2009. ISSN 1557-7988 (Electronic) 0076-6879 (Linking). Disponível em: < https://www.ncbi.nlm.nih.gov/pubmed/19903557 >.

Rohou, A.; Grigorieff, N. CTFFIND4: Fast and accurate defocus estimation from electron micrographs. J Struct Biol, v. 192, n. 2, p. 216–21, Nov 2015. ISSN 1095-8657 (Electronic) 1047-8477 (Linking). Disponível em: < https://www.ncbi.nlm.nih.gov/pubmed/26278980 >.

Ruan, Y. et al. Direct visualization of glutamate transporter elevator mechanism by highspeed AFM. Proceedings of the National Academy of Sciences of the United States of America, v. 114, n. 7, p. 1584–1588, Feb 14 2017. ISSN 0027-8424. Disponível em: < <Go to ISI>://WOS:000393989300061 >.

Rusinova, R. et al. Regulation of ion channel function by the host lipid bilayer examined by a stopped-flow spectrofluorometric assay. Biophys J, v. 106, n. 5, p. 1070–8, Mar 4 2014. ISSN 1542-0086 (Electronic) 0006-3495 (Linking). Disponível em: < https://www.ncbi.nlm.nih.gov/pubmed/24606931 >.

Scheres, S. H. Processing of Structurally Heterogeneous Cryo-EM Data in RELION. Methods Enzymol, v. 579, p. 125–57, 2016. ISSN 1557-7988 (Electronic) 0076-6879 (Linking). Disponível em: < https://www.ncbi.nlm.nih.gov/pubmed/27572726 >.

Scopelliti, A. J. et al. Structural characterisation reveals insights into substrate recognition by the glutamine transporter ASCT2/SLC1A5. Nature Communications, v. 9, Jan 2 2018. ISSN 2041-1723. Disponível em: < <Go to ISI>://WOS:000419306000029 >.

Tzingounis, A. V. et al. Arachidonic acid activates a proton current in the rat glutamate transporter EAAT4. J Biol Chem, v. 273, n. 28, p. 17315–7, Jul 10 1998. ISSN 0021-9258 (Print) 0021-9258 (Linking). Disponível em: < https://www.ncbi.nlm.nih.gov/pubmed/9651313 >.

Verdon, G.; Boudker, O. Crystal structure of an asymmetric trimer of a bacterial glutamate transporter homolog. Nature Structural & Molecular Biology, v. 19, n. 3, p. 355–357, Mar 2012. ISSN 1545-9993. Disponível em: < <Go to ISI>://WOS:000301181900013 >.

Verdon, G. et al. Coupled ion binding and structural transitions along the transport cycle of glutamate transporters. Elife, v. 3, May 19 2014. ISSN 2050-084x. Disponível em: < <Go to ISI>://WOS:000336040700004 >.

Voss, N. R. et al. DoG Picker and TiltPicker: software tools to facilitate particle selection in single particle electron microscopy. J Struct Biol, v. 166, n. 2, p. 205–13, May 2009. ISSN 1095-8657 (Electronic) 1047-8477 (Linking). Disponível em: < https://www.ncbi.nlm.nih.gov/pubmed/19374019 >.

Yernool, D. et al. Structure of a glutamate transporter homologue from Pyrococcus horikoshii. Nature, v. 431, n. 7010, p. 811–818, Oct 14 2004. ISSN 0028-0836. Disponível em: < <Go to ISI>://WOS:000224435500038 >.

Yu, X. D. et al. Cryo-EM structures of the human glutamine transporter SLC1A5 (ASCT2) in the outward-facing conformation. Elife, v. 8, Oct 3 2019. ISSN 2050-084x. Disponível em: < <Go to ISI>://WOS:000491168300001 >.

Zerangue, N. et al. Differential modulation of human glutamate transporter subtypes by arachidonic acid. J Biol Chem, v. 270, n. 12, p. 6433–5, Mar 24 1995. ISSN 0021-9258 (Print) 0021-9258 (Linking). Disponível em: < https://www.ncbi.nlm.nih.gov/pubmed/7896776 >.

Zheng, S. Q. et al. MotionCor2: anisotropic correction of beam-induced motion for improved cryo-electron microscopy. Nat Methods, v. 14, n. 4, p. 331–332, Apr 2017. ISSN 1548-7105 (Electronic) 1548-7091 (Linking). Disponível em: < https://www.ncbi.nlm.nih.gov/pubmed/28250466 >.

Zhou, W. et al. Large-scale state-dependent membrane remodeling by a transporter protein. Elife, v. 8, Dec 19 2019. ISSN 2050-084X (Electronic) 2050-084X (Linking). Disponível em: < https://www.ncbi.nlm.nih.gov/pubmed/31855177 >.

Zivanov, J. et al. New tools for automated high-resolution cryo-EM structure determination in RELION-3. Elife, v. 7, Nov 9 2018. ISSN 2050-084X (Electronic) 2050-084X (Linking). Disponível em: < https://www.ncbi.nlm.nih.gov/pubmed/30412051 >.

